# Small proteome of the nitrogen-fixing plant symbiont *Sinorhizobium meliloti*

**DOI:** 10.1101/2022.11.12.516264

**Authors:** Lydia Hadjeras, Benjamin Heiniger, Sandra Maaß, Robina Scheuer, Rick Gelhausen, Saina Azarderakhsh, Susanne Barth-Weber, Rolf Backofen, Dörte Becher, Christian H. Ahrens, Cynthia M. Sharma, Elena Evguenieva-Hackenberg

## Abstract

The soil-dwelling plant symbiont *Sinorhizobium meliloti* is a major model organism of Alphaproteobacteria. Despite numerous detailed OMICS studies, information about small open reading frame (sORF)-encoded proteins (SEPs) is largely missing, because sORFs are poorly annotated, and SEPs are hard to detect experimentally. However, given that SEPs can fulfill important functions, cataloging the full complement of translated sORFs is critical for analyzing their roles in bacterial physiology. Ribosome profiling (Ribo-seq) can detect translated sORFs with high sensitivity, but is not yet routinely applied to bacteria because it must be adapted for each species. Here, we established a Ribo-seq procedure for *S. meliloti* 2011 based on RNase I digestion and detected translation for 60% of the annotated coding sequences during growth in minimal medium. Using ORF prediction tools based on Ribo-seq data, subsequent filtering, and manual curation, the translation of 37 non-annotated sORFs with ≤ 70 amino acids was predicted with high confidence. The Ribo-seq data were supplemented by mass spectrometry (MS) analyses from three sample preparation approaches and two integrated proteogenomic search databases (iPtgxDBs). Searches against a standard and a 20-fold smaller Ribo-seq data-informed custom iPtgxDB confirmed many annotated SEPs and identified 11 additional novel SEPs. Epitope tagging and Western blot analysis confirmed the translation of 15 out of 20 SEPs selected from the translatome map. Overall, by applying MS and Ribo-seq as complementary approaches, the small proteome of *S. meliloti* was substantially expanded by 48 novel SEPs. Several of them are conserved from *Rhizobiaceae* to Bacteria, suggesting important physiological functions.

## INTRODUCTION

Over the last two decades, using next-generation sequencing and high throughput OMICS profiling technologies, the genomes of thousands of bacteria have been assembled, and the transcriptomes and proteomes of many of them have been analyzed under different conditions, with the aim of gaining insights into the genetic and molecular basis of their biology. Despite this wealth of data, information about small open reading frame (sORF)-encoded proteins (SEPs), which are proteins with less than 50 or 100 amino acids (aa), is scarce (Duval and Cossart 2017; Gray et al. 2022; Hemm et al. 2020; Orr et al. 2020; Storz et al. 2014). Recently, the small proteomes of eukaryotes, bacteria, and viruses have been focused on, as a growing number of small proteins have been demonstrated to fulfill important physiological functions, such as in cell division, metabolism, transport, signal transduction, spore formation, cell communication, cellular stress responses, and virulence (Aoyama et al. 2022; Duval and Cossart 2017; Hemm et al. 2020; Melior et al. 2020; Khitun and Slavoff 2019; Patraquim et al. 2020; Song et al. 2022; Storz et al. 2014). Therefore, cataloging the full complement of small proteins is critical in achieving a more comprehensive and accurate description of the proteomes of bacterial model organisms and their potential functions.

Small protein identification is difficult due to several technical challenges. For instance, SEPs are difficult to detect using SDS-PAGE or mass spectrometry (MS) for various technical reasons (Ahrens et al. 2022; Fijalkowski et al. 2022; Storz et al. 2014). Limitations of standard shotgun proteomics workflows at the sample preparation, protease digestion, liquid chromatography, MS data acquisition, and bioinformatic data analysis steps affect comprehensive MS-based SEP identification (Ahrens et al. 2022; Cassidy et al. 2021). Furthermore, variable length thresholds were used in the genome annotation step to minimize the number of spurious ORF predictions. As a result, sORFs encoding truly expressed small proteins are often missing from genome annotations (Hahn et al. 2016; Storz et al. 2014). Meanwhile, various strategies to achieve extensive proteome coverage of the notoriously under-represented classes of small and membrane proteins (novel small proteins are often membrane associated) have been applied for prokaryotes (Omasits et al. 2013; Wiśniewski 2016; Zhang et al. 2013). Methods to enrich bacterial SEPs in samples are further improved, for example, with the use of small pore-sized solid-phase materials ( Bartel et al. 2020; Cassidy et al. 2019; Petruschke et al. 2020), and digestion with alternative/multiple proteases has been performed to increase the number of identified SEPs ( Bartel et al. 2020; Kaulich et al. 2021; Petruschke et al. 2021). The obtained mass spectra are usually assigned to peptide or protein sequences by matching the determined fragment ion masses to the predictions derived from a sequence database (DB). Therefore, only peptides with sequences available in the protein search DB can be identified. Consequently, custom protein search DBs that try to capture the entire coding potential of prokaryotic genomes have been proposed, such as integrated proteogenomic search DBs (iPtgxDBs). They integrate and consolidate the differences among existing reference genome annotations, *ab initio* gene predictions, and a modified six-frame translation by considering alternative start sites, thereby enabling the detection of novel proteins, including SEPs (Omasits et al. 2017). Thus, proteogenomic studies that combine results from SEP-optimized MS data searched with iPtgxDBs or other custom search DBs and ribosome profiling (Ribo-seq) have great potential to detect highly comprehensive and accurate compendia of novel small proteins.

Ribo-seq has developed into a powerful method to study and annotate translatome globally, including sORFs (Ingolia 2016; Vazquez-Laslop et al. 2022). Compared with MS-based proteomics, Ribo-seq has the advantage of higher sensitivity (Ahrens et al. 2022; Duval and Cossart 2017; Gray et al. 2022; Hemm et al. 2020; Orr et al. 2020; Venturini et al. 2020; Storz et al. 2014). Ribo-seq relies on deep sequencing of approximately 30-nt-long “footprint” regions of the mRNA bound by the ribosome during translation and protected against nuclease digestion. In addition to providing a global picture of actively translated mRNAs in the cell, Ribo-seq also reveals the specific location on the RNA where the ribosome was bound, allowing the mapping of ORFs. For this, cells are lysed under certain conditions, allowing for the “freezing” of ribosomes on mRNAs. mRNA parts that are not protected by the ribosomes are then digested to generate ribosome footprints that are sequenced and mapped to the genome (Ingolia et al. 2009; Ingolia 2016). While Ribo-seq-based detection of translated mRNA works well for eukaryotic cells at a single codon resolution, this method is difficult to utilize for prokaryotes (Glaub et al. 2020; Mohammad et al. 2019; Vazquez-Laslop et al. 2022). Nevertheless, adapting and refining the Ribo-seq method has enabled the detection of many new, translated sORFs and corresponding SEPs not only in *Escherichia coli* but also in several other bacterial species and in halophilic archaea (Gelsinger et al. 2020; Meydan et al. 2019; Mohammad et al. 2019; Vazquez-Laslop et al. 2022; Weaver et al. 2019). However, for many bacterial model organisms, Ribo-seq data are still missing, as the protocols typically have to be adapted and optimized for each bacterial organism (Duval and Cossart 2017; Gray et al. 2022; Hemm et al. 2020; Orr et al. 2020; Venturini et al. 2020; Storz et al. 2014).

*Sinorhizobium meliloti* is an agriculturally important bacterial species that lives in soil and can fix molecular nitrogen in symbiosis with legume plants (Jones et al. 2007). Due to its versatile lifestyle and ecological relevance, it is a major model organism for studying gene regulation in Alphaproteobacteria. In addition, its relatively close relationship to pathogens of the genus *Brucella* makes *S. meliloti* an attractive model for host–pathogen research (Marlow et al. 2009). Several OMICS datasets are available for *S. meliloti* 2011 and its sibling, *S. meliloti* 1021, which is the first sequenced strain of this species (Galibert et al. 2001). These consist of proteomics (Djordjevic 2004; Barra-Bily et al. 2010; Sobrero et al. 2012; Marx et al. 2016) and transcriptomic datasets, including differential RNA-seq that enables the annotation of transcription start sites, 5′ and 3′ UTRs, and novel transcripts (Becker et al. 2004; Schlüter et al. 2013; Sallet et al. 2013). The *S. meliloti* 2011 6.7 Mb genome harbors a 3.66 Mb chromosome and two megaplasmids, the 1.35 Mb pSymA and the 1.68 Mb pSymB (Sallet et al. 2013). As a proof of principle, an iPtgxDB created for *S. meliloti* 2011 has allowed the detection of the 14-aa-long leader peptide peTrpL in the proteomic data, in which a function in resistance to multiple antimicrobial compounds can be subsequently established (Melior et al. 2020). However, the identification of additional functional SEPs in *S. meliloti* and related Alphaproteobacteria has been limited by the lack of studies specifically targeting the small proteome and translatome.

Here, we developed and then applied Ribo-seq on *S. meliloti* 2011 to map its translatome globally, with a focus on the small proteome (data available at our interactive web-based genome-browser: http://www.bioinf.uni-freiburg.de/ribobase). The use of RNase I in our Ribo-seq showed successful trimming of mRNA regions that were not protected by ribosomes, allowing differentiation between translated and untranslated regions. Besides detecting the translation of annotated sORFs (some of which are available in recent updates of the genome annotation), we also uncovered 37 translated novel, non-annotated sORFs located on different replicons. The translation of several annotated, as well as novel, sORFs was further validated by MS-based proteomics using iPtgxDBs and/or epitope tagging and Western blot analysis, thereby confirming predictions based on Ribo-seq coverage. Eleven novel SEPs were uniquely identified by MS, showing that using both methods when mapping the small proteome is advantageous. Overall, our combined approach provided a set of 48 novel *S. meliloti* sORFs, many of which are conserved, as a resource to further elucidate their roles in bacterial physiology and symbiosis.

## METHODS

### Growth and harvest of *S. meliloti* for Ribo-seq

*S. meliloti* 2011 (Casse et al. 1979) was first cultivated on TY (5 g of BactoTryptone, 3 g of Bacto-yeast extract, and 0.3 g of CaC_2_ per liter) agar plates (Beringer 1974). The plate cultures were used to inoculate liquid cultures, which were grown semi-aerobically (routinely, 30 ml of medium in a 50 ml Erlenmeyer flask under constant agitation at 140 rpm) at 30 °C in GMS minimal medium (10 g of D-mannitol, 5 g of sodium glutamate, 5 g of K_2_HPO_4_, 0.2 g of MgSO_4_ × 7H_2_O, and 0.04 g of CaCl_2_ per liter; trace elements: 0.05 mg of FeCl_3_ × 6H_2_O, 0.01 mg of H_3_Bo_3_, 0.01 mg of ZnSO_4_ × 7H_2_O, 0.01 mg of CoCl_2_ × 6H_2_O, 0.01 mg of CuSO_4_ × 5H_2_O, 1.35 mg of MnCl_2_, and 0.01 mg of Na_2_MoO_4_ × 2H_2_O per liter; 10 μg of biotin and 10 mg of thiamine per liter) ^(^Zevenhuizen and van Neerven 1983). As the strain exhibits chromosomally encoded streptomycin resistance, 250 µg/ml streptomycin was added to the media. For Ribo-seq sample preparation, cells corresponding to 40 OD_600_ equivalent units were harvested after rapid chilling in an ice bath to halt cell growth and translation. In brief, cultures in the exponential phase (OD_600nm_ 0.5) were rapidly placed in a pre-chilled flask in an ice-water bath and incubated with gentle shaking for 3 min. Cells were then immediately pelleted by centrifugation (10 min at 6000 ×g) before snap-freezing in liquid N_2_. Before centrifugation, a culture aliquot was withdrawn for total RNA analysis, mixed with 0.2 vol stop mix (5% buffer-saturated phenol [Roth] in 95% ethanol), and snap-frozen in liquid N_2_. Even though translation elongation inhibitors have been extensively used in both eukaryotic and bacterial Ribo-seq workflows, using such chemicals can introduce bias into Ribo-seq coverage (Gerashchenko and Gladyshev 2014; Mohammad et al. 2019). Therefore, we chose to perform Ribo-seq without these inhibitors because we were able to recover sufficient polysomes using the fast-chilling method (see **Fig. 1**).

### Preparation of ribosome footprints

Ribo-seq was performed as previously described (Oh et al. 2011), with some modifications. In brief, cell pellets were resuspended with cold lysis buffer (1 M NH_4_Cl, 150 mM MgCl_2_, 20 mM Tris-HCl, 5 mM CaCl_2_, 0.4% Triton X-100, 150 U DNase I [Fermentas], and 1000 U RNase Inhibitor [MoloX, Berlin] at pH 8.0) and lysed by sonication (constant power 50%, duty cycle 50%, and 3 × 30 s cycles with 30 s cooling on a water-ice bath between each sonication cycle to avoid heating of the sample). The lysate was clarified by centrifugation at 10,000 × g for 12 min at 4 °C. Approximately 15 A_260_ of lysate 200 U of RNase I (Thermo Fisher Scientific) was added. Polysome digestion was performed at 25°C with shaking at 650 rpm for 90 min. A mock-digested control (no enzyme added) was performed in parallel to confirm the presence of polysomes in the lysate. To analyze polysome profiles and recover digested monosomes, we layered 15 A_260_ units onto a linear 10%–55% sucrose gradient prepared in 4X gradient buffer (10X gradient buffer: 100 mM MgCl_2_, 200 mM Tris-HCl, 1 M NH_4_Cl, and 20 mM dithiothreitol [DTT] at pH 8.0) in an ultracentrifuge tube (13.2 mL Beckman Coulter SW-41). Gradients were centrifuged in a SW40-Ti rotor at 35,000 rpm for 2 h and 30 min at 4 °C in a Beckman Coulter Optima XPN-80 ultracentrifuge. Gradients were processed using a gradient station (IP, Biocomp Instruments) fractionation system with continuous absorbance monitoring at 254 nm to resolve ribosomal subunit peaks. The 70S monosome fractions were collected and subjected to RNA extraction to purify the RNA footprints. RNA was extracted from fractions or cell pellets for total RNA using hot phenol-chloroform-isoamyl alcohol (25:24:1, Roth) or hot phenol (Roth), respectively, as previously described (Sharma et al. 2007; Venturini et al. 2020). Ribosomal RNA (rRNA) was depleted from 5 µg of DNase I-digested total RNA by subtractive hybridization with the Pan-Bacteria riboPOOLs (siTOOLs, Germany) according to the manufacturer’s protocol with Dynabeads MyOne Streptavidin T1 beads (Invitrogen). Total RNA was fragmented with an RNA fragmentation reagent (Ambion). Monosome RNA and fragmented total RNA were size selected (26–34 nt) on 15% polyacrylamide/7 M urea gels, as previously described (Ingolia et al. 2012) using RNA oligonucleotides NI-19 and NI-20 as guides. RNA was cleaned and concentrated by isopropanol precipitation with 15 μg of GlycoBlue (Ambion) and dissolved in H_2_O. cDNA libraries were prepared by Vertis Biotechnologie AG (Freising, Germany) using the adapter ligation protocol without fragmentation. First, an oligonucleotide adapter was ligated to the 3′ end of the RNA molecules. First-strand cDNA synthesis was performed using M-MLV reverse transcriptase and the 3′ adapter as the primer. The first strand of cDNA was purified, and the 5′ Illumina TruSeq sequencing adapter was ligated to the 3′ end of the antisense cDNA. The resulting cDNA was PCR-amplified to approximately 10–20 ng/μl using a high-fidelity DNA polymerase. The DNA was purified using the Agencourt AMPure XP kit (Beckman Coulter Genomics) and analyzed by capillary electrophoresis. The primers used for PCR amplification were designed for TruSeq sequencing according to the instructions of Illumina. The following adapter sequences flank the cDNA inserts: TruSeq_Sense_primer: (NNNNNNNN = i5 Barcode for multiplexing) 5′-AATGATACGGCGACCACCGAGATCTACAC-NNNNNNNN-ACACTCTTTCCCTACA CGACGCTCTTCCGATCT-3′; TruSeq_Antisense_primer: (NNNNNNNN = i7 Barcode for multiplexing) 5′-CAAGCAGAAGACGGCATACGAGAT-NNNNNNNN-GTGACTGGAGTTCAGACGTGT GCTCTTCCGATCT-3′. cDNA libraries were pooled on an Illumina NextSeq 500 high-output flow cell and sequenced in single-end mode (75 cycles, with 20 million reads per library) at the Core Unit SysMed at the University of Würzburg.

### Ribo-seq data analysis

*S. meliloti* Ribo-seq data were processed and analyzed using the published HRIBO workflow (version 1.6.0) (Gelhausen et al. 2021), which has previously been used for the analysis of bacterial Ribo-seq data (Venturini et al. 2020). In brief, sequencing read files were processed with a snakemake (Köster and Rahmann 2012) workflow, which downloads all required tools from bioconda (Grüning et al. 2018) and automatically determines the necessary processing steps. Adapters were trimmed from the reads with cutadapt (version 2.1) (Martin 2011) and then mapped against the *S. meliloti* 2011 genome with segemehl (version 0.3.4) (Otto et al. 2014). Reads corresponding to rRNA and other multiply mapping reads were removed with SAMtools (version 1.9) (Li et al. 2009). Quality control was performed by creating read count statistics for each processing step and RNA class with Subread featureCounts (1.6.3) (Liao et al. 2014). All processing steps were analyzed with FastQC (version 0.11.8) (Wingett and Andrews 2018), and the results were aggregated with MultiQC (version 1.7) (Ewels et al. 2016). Summary statistics are shown in **Table S1**.

Read coverage files were generated with HRIBO using different full-read mapping approaches (global or centered) and single-nucleotide mapping strategies (5′ or 3′ end). Read coverage files using two different normalization methods were created (mil and min). For the mil normalization, read counts were normalized by the total number of mapped reads within the sample and scaled by a per-million factor. For the min normalization, the read counts were normalized by the total number of mapped reads within the sample and scaled by the minimum number of mapped reads among all analyzed samples. The coverage files generated using the min normalization and the global mapping (full read) approach were used for genome browser visualization. Metagene analysis of ribosome density at start codons was performed as previously described (Becker et al. 2013).

### Ribo-seq-based ORF prediction, filtering, and manual curation

ORFs were called with an adapted variant of REPARATION (Ndah et al. 2017) using blast instead of usearch (see https://github.com/RickGelhausen/REPARATION_blast) and DeepRibo (Clauwaert et al. 2019). In addition, generic feature format (GFF) track files with the same information were created for in-depth manual genome browser inspection, as well as GFF files showing potential start and stop codons and ribosome binding site information. Summary statistics for all available GenBank annotated and merged novel ORFs detected by REPARATION and DeepRibo were computed in a tabularized form, including, among other values, translation efficiency (TE), RPKM (reads per kilobase of transcript per million mapped reads) normalized read counts, codon counts, and nucleotide and aa sequences (see **Table S2**). Annotated sORFs were classified as translated if they fulfilled an arbitrary mean TE cut-off of ≥ 0.5 and RNA-seq and Ribo-seq RPKM of ≥ 10 (cut-offs chosen based on the lowest TE and RPKM values associated with housekeeping genes [i.e., ribosomal protein genes] and the genes detected by proteomics). To identify strong candidates for novel sORFs, we inspected HRIBO ORF predictions from DeepRibo and REPARATION. As DeepRibo is prone to a high rate of false positives (Gelhausen et al. 2022), we first generated a reasonable set of potential novel sORFs by applying the following expression cut-off filters: mean TE of ≥ 0.5 and RNA-seq and Ribo-seq RPKM of ≥10 (in both replicates) based on the 85 positively labeled translated sORFs (see **Fig. 3**). In addition, novel translated sORF candidates were required to be predicted by DeepRibo with a prediction score of > −0.5 that allows for ORF candidate ranking (Clauwaert et al. 2019). The filtered sORFs were then subjected to manual curation, that is, inspection of the Ribo-seq coverage in a genome browser. To assert translation, we considered the evenness of the Ribo-seq coverage, its restriction to the predicted ORF, and the high read coverage in the footprint library compared with the corresponding transcriptome library. We created an interactive web-based genome browser using JBrowse (http://www.bioinf.uni-freiburg.de/ribobase) (Buels et al. 2016), where the coverage files for the Ribo-seq replicates, the annotation, and the predicted sORF can be visualized.

### Sample preparation for MS

For MS analysis, cells of 1.5 l of an *S. meliloti* culture (OD_600nm_ 0.5) were harvested by centrifugation at 6,000 rpm and 4°C. The cell pellet was resuspended in 30 ml of buffer containing 20 mM Tris, 150 mM KCl, 1 mM MgCl2, and 1 mM DTT at pH 7.5. After lysis by sonication and centrifugation at 13,000 rpm for 30 min at 4°C, the cleared lysates were frozen in liquid nitrogen and stored at −80 °C. To generate a highly comprehensive small protein dataset, we used three complementary approaches for sample preparation: 1) tryptic in-solution digest of all proteins in the sample, 2) solid-phase enrichment (SPE) of small proteins without any subsequent digestion, and 3) SPE of small proteins with subsequent digestion using Lys-C. Sample preparation was performed as previously described (Bartel et al. 2020) with some modifications. In brief, samples for tryptic in-solution digests were reduced and alkylated before trypsin was added in an enzyme-to-protein ratio of 1:100, and samples were incubated at 37 °C for 14 h. The digest was stopped by acidifying the mixture with HCl. For SPE, samples were loaded on an equilibrated column packed with an 8.5 nm pore size, modified styrene-divinylbenzene resin (8B-S100-AAK, Phenomenex), which was then washed to remove large proteins. The enriched small protein fraction was eluted with 70% (v/v) acetonitrile and evaporated to dryness in a vacuum centrifuge. The SPE samples were either directly used for MS or in-solution digested as described above but with Lys-C instead of trypsin.

### Generation of standard and custom iPtgxDBs to identify novel SEPs

iPtgxDBs were generated based on the *S. meliloti* 2011 ASM34606v1 reference genome sequence as described (Omasits et al. 2017). Annotations from several reference genome centers and/or releases (GenBank 2014, RefSeq2017, Genoscope), two *ab initio* gene predictions (Prodigal, ChemGenome), and *in silico* ORF predictions were hierarchically integrated for a trypsin-specific iPtgxDB as previously detailed (Melior et al. 2020), (https://iptgxdb.expasy.org/database/annotations/s-meliloti-tryptic; see **Table S3.1**). To capture data from all three experimental approaches, two more iPtgxDBs were created in a similar fashion using command-line utilities. While default settings were used to create the trypsin-specific iPtgxDB, for the LysC-specific iPtgxDB, the regular expression “(K)(.)” was used, allowing cleavage after every lysine. The iPtgxDB for the experiments without protease digestion was generated with a regular expression that did not allow any cleavages. In addition, three 20-fold smaller custom iPtgxDBs were created to improve search statistics/predictive potential. For these, instead of adding the Chemgenome and *in silico* predictions, 266 selected Ribo-seq translation products identified from the sORF prediction tools DeepRibo (Clauwaert et al. 2019) and Reparation (Ndah et al. 2017), as well as manual analysis, were converted to GFF format using a custom Python script and integrated along with the RefSeq, GenBank, Genoscope, and Prodigal predictions to create the respective iPtgxDBs (**Tables S3.3** and **S3.4**). All six iPtgxDBs (downloadable from https://iptgxdb.expasy.org) also contained sequences of common laboratory contaminants (116 from CrapOme and 256 from the Functional Genomics Center Zurich). All peptides implying potentially novel proteins were subjected to a PeptideClassifier analysis (Qeli and Ahrens 2010) extended for proteogenomics in prokaryotes (Omasits et al. 2017). This procedure ensures that i) only unambiguous peptides were considered (class 1a) or ii) annotation cluster-specific cases can be distinguished: Class 2a peptides imply a subset of all possible proteoforms (e.g., like an extension, reduction), class 2b peptides imply all isoforms, which means that the gene encoding the proteoforms, but not a specific proteoform, was identified.

### MS analysis

Samples were loaded on an EASY-nLC 1200 (Thermo-Fisher Scientific) equipped with an in-house-built 20 cm reversed-phase column packed with 3 µm Reprosil-Pur 120 C18-AQ (Dr. Maisch) and an integrated emitter tip. Peptides were eluted by a 156 min non-linear gradient of solvent B (0.1% v/v acetic acid in acetonitrile) and injected online in an Orbitrap Velos (Thermo-Fisher Scientific). The survey scans were acquired in the Orbitrap (300–1700 Th; 60,000 resolution at 400 m/z; 1 × 1e^6^ predictive automatic gain control target; activated lock mass correction). After collision-induced dissociation with a normalized collision energy of 35, fragment spectra were recorded in the LTQ (mass range dependent on precursor m/z; 3 × 1e^4^ predictive automatic gain control) for the 20 most abundant ions. Fragmented ions were dynamically excluded from fragmentation for 30 s.

DB searches were performed with Sorcerer-SEQUEST 4 (Sage-N Research, Milpitas, USA), allowing two missed cleavages for samples derived from tryptic in solution digest or LysC digested SPE samples and with non-specified enzymes for SPE samples without proteolytic digest. No fixed modifications were considered, and oxidation of methionine was considered a variable modification. The mass tolerance for precursor ions was set to 10 ppm, and the mass tolerance for fragment ions was set to 1.0 Da. Validation of MS/MS-based peptide and protein identification was performed with Scaffold V4.8.7 (Proteome Software, Portland, USA), and peptide identifications were accepted if they exhibited at least deltaCn scores of > 0.1 and XCorr scores of > 2.2, 3.3, and 3.75 for doubly, triply, and all high-charged peptides, respectively. Identifications for proteins of > 15 kDa were only accepted if at least two unique peptides were identified. Proteins that contained ambiguous, non-unique peptides and could not be differentiated based on MS/MS analysis alone were grouped to satisfy the principles of parsimony (Sorcerer-SEQUEST). Identifications for annotated proteins of < 15 kDa were accepted if at least one unique peptide was identified with at least two peptide spectrum matches (PSMs). To identify novel proteins, we required additional PSM evidence from predictions as described before (Varadarajan et al. 2020a, 2020b), that is, 3 PSMs for *ab initio* predictions and 4 PSMs from *in silico* predictions. Here, we also allowed *in silico* candidates with 3 PSMs if they were observed in each of the three replicates. Similar to the RefSeq annotated proteins, novel proteins longer than 150 aa required two unique peptides (however, these were not the focus of this study). The application of these filter criteria kept the protein false discovery rate (FDR) below 1%. To facilitate overview and comparison, we integrated MS-identified proteins, Ribo-Seq, and Western blot analysis data in a “master table” (**Table S4**).

### Cloning procedures

The oligonucleotides (Microsynth) used for cloning are listed in **Table S5**. Routinely, FastDigest Restriction Endonucleases and Phusion polymerase (Thermo Fisher Scientific) were used. PCR products were first ligated into pJet1.2/blunt (CloneJet PCR Cloning Kit, Thermo Fisher Scientific) and transformed into *E. coli* DH5-alpha. Subsequently, inserts were subcloned in conjugative plasmids originating from pSRKGm (Khan et al. 2008). Insert sequences were analyzed by Sanger sequencing with plasmid-specific primers (Microsynth Seqlab). *E. coli* S17-1 was used to transfer the plasmids to *S. meliloti* 2011 by diparental conjugation (Simon et al. 1983).

Plasmid pSW2 was used to clone the candidate sORFs. It was constructed using pRS1, a derivative of pSRKGm, in which the *E. coli lac* module was exchanged for a multiple cleavage site-containing cloning site for the restriction endonucleases NheI, HindIII, XbaI, SpeI, BamHI, PstI, and EcoRI. First, a transcription terminator T_*rrn*_ from *Bradyrhizobium japonicum* USDA 110 was cloned into the EcoRI restriction site of pRS1. For this, the terminator containing sequence was amplified with the forward primer Bj-Trrn-Fw-2019 and the reverse primer Bj-Trrn-Rv-2019 using plasmid pJH-O1 as a template (Čuklina et al. 2016). In the PCR product, an EcoRI restriction site was present downstream of the forward primer sequence. This restriction site and that in the reverse primer were used for the transcription terminator cloning. A clone with the desired orientation was selected, and the plasmid was named pRS1-Trrn (**Fig. S1**). Double-stranded DNA encoding a sequential peptide affinity (SPA) tag, which is composed of the calmodulin-binding peptide and three modified FLAG sequences separated by a TEV protease cleavage site (Zeghouf et al. 2004), was then cloned between the BamHI and EcoRI cleavage sites of pRS1-Trrn. The SPA-tag encoding sequence was designed without an ATG codon, without rare codons, and with Gly-Gly-Gly-Ser linker codons at the 5′ end and adapted to the high GC content of *S. meliloti*. It was generated synthetically by Eurofins and provided on plasmid pEX-A128, which was used as a template for PCR amplification with primers SmSPA-Ct-BamFW and SmSPA-Ct-EcoRv. The resulting plasmid pSW1 can be used to clone an sORF in frame with the SPA-encoding sequence and under the control of its own promoter. Here, pSW1 was used to clone the promoter P*sinI* between the NheI and XbaI restriction sites. The promoter sequence (McIntosh et al. 2008) was amplified using primers NheI-PsinI-FW and XbaI-PsinI-RV and *S. meliloti* 2011 genomic DNA as a template. The resulting pSW2 plasmid was used to clone candidate sORFs, each with a 15-nt upstream region potentially harboring a Shine-Dalgarno sequence between the XbaI and BamHI restriction sites (**Fig. S1**). In total, 20 sORF::SPA fusions were cloned and tested by Western blot analysis. The corresponding plasmids were designated from pSW2-SEP1 to pSW2-SEP20.

### Western blot analysis

Exponentially grown *S. meliloti* cells (OD_600nm_ 0.5; minimal medium) were harvested (3,500 ×g for 10 min at 4 °C) and resuspended in an SDS-loading buffer. After incubation for 5 min at 95 °C, the crude lysate proteins were separated by Tricine-SDS PAGE (16%) and blotted onto a PVDF membrane (Amerham^TM^Hybond^TM^, 0.2 µM PVDF; GE Healthcare Life Science, Chalfont St Giles, Great Britain) as described (Schägger 2006). For detection, monoclonal ANTI-FLAG M2-Peroxidase (HRP) antibodies (Merck, Darmstadt, Germany) and Lumi-Light Western-Blot-Substrate (Roche, Basel, Schweiz) were used. Signal visualization was performed with a chemiluminescence imager (Fusion SL4, Vilber, Eberhardzell, Germany). For fractionation, the cell pellets were resuspended in TKMDP buffer containing 20 mM Tris-HCl, 150 mM KCl, 1 mM MgCl_2_, 1 mM DTT, and one protease inhibitor cocktail tablet at pH 7.5 (Sigma Aldrich, St. Louis, USA). Lysates prepared by three passages in a French press at 1,000 psi were cleared by centrifugation at 14,000 ×g for 30 min at 4 °C. The supernatant was subjected to ultracentrifugation at 100,000 ×g for 1 h at 4 °C. The supernatant (S100 fraction) was then removed, and the P100 pellet was resuspended in the same volume of TKMDP buffer.

### Conservation and domain search analyses

The identification of novel small protein homologues was performed using Blastp and tBlastn searches in bacteria on the National Center for Biotechnology Information DB (https://blast.ncbi.nlm.nih.gov/Blast.cgi). The protein sequences for novel protein candidates identified by Ribo-seq and/or MS were used as query sequences. For tBlastn, the following parameters were used: the filter for low complexity regions off, a seed length that initiates an alignment (word size) of 6, 60% coverage of the query sequence with at least 40% identity, an E-value (Expect value) of ≤100 to capture all potential orthologs, and an E-value between 0.01 and 1 for high-confidence hits (Allen et al. 2014). Moreover, novel small proteins discovered in this study were further analyzed for secondary structure and predicted protein domains and predictions of lipoproteins, as well as potential subcellular localization using predictions from the Phyre2 v2.0 (http://www.sbg.bio.ic.ac.uk/∼phyre2/), LipoP-1.0 (https://services.healthtech.dtu.dk/service.php?LipoP-1.0) TMHMM v2.0 (https://services.healthtech.dtu.dk/service.php?TMHMM-2.0), and PSORTb v3.0.2 servers (https://www.psort.org/psortb/).

### Data availability

The MS-based proteomics data were deposited to the ProteomeXchange Consortium at the PRIDE partner repository, with dataset identifier PXD034931. The iPtgxDBs can be downloaded from https://iptgxdb.expasy.org/. Ribo-seq and RNA-seq data were deposited in GEO, with accession number GSE206492. The Ribo-seq and RNA-seq data of *S. meliloti* 2011 can be viewed with an interactive online JBrowse instance (http://www.bioinf.uni-freiburg.de/ribobase).

## RESULTS

### Establishing Ribo-seq in *S. meliloti* to map its translatome

To provide a genome-wide map of translated annotated sORFs and to reveal new sORFs in the plant symbiont *S. meliloti*, we first adapted the Ribo-seq protocol (Oh et al. 2011) to this organism (**Fig. 1A**). For this purpose, several steps, including cell harvest, lysis, and footprint generation, were optimized (see Methods). *S. meliloti* 2011 cells were grown to the mid-log phase in minimal medium, and samples were rapidly cooled and harvested to avoid polysome run-off. Polysome profile analysis after lysate fractionation on a sucrose gradient showed successfully captured translating ribosomes (**Fig. 1B**, black profile). The mRNA should be ribonucleolytically digested outside ribosomes to produce ribosome footprints. Since the broad-range ribonuclease RNase I, which is often used for eukaryotic Ribo-seq analysis, is inactive on polysomes from enteric bacteria (Bartholomäus et al. 2016), most prokaryotic Ribo-seq protocols mainly use micrococcal nuclease (MNase) instead. However, MNase preferentially cleaves at pyrimidines, introduces periodicity artifacts, and generates footprints that are more heterogeneous in length than those from RNase I (Ingolia 2016; Vazquez-Laslop et al. 2022). Therefore, we used RNase I to convert *S. meliloti* polysomes into monosomes (**Fig. 1B**) and to generate ribosome footprints (**Fig. 1C–1E**). By comparing Ribo-seq read coverage data and expression signals from a paired RNA-seq library generated from fragmented total RNA, features, such as coding potential, ORF boundaries, and 5′ and 3′ UTRs, can be defined (**Fig. 1C–1E**).

Inspection of Ribo-seq coverage for translated ORFs and known non-coding transcripts further demonstrated the successful setup of Ribo-seq in *S. meliloti.* For example, the protein-coding genes *rpsO* and *icd* showed higher cDNA read coverage in the Ribo-seq library compared with the paired RNA-seq library (**Fig. 1C**), whereas the RNase P RNA gene *rnpB* showed high cDNA read coverage only in the RNA-seq library (**Fig. 1D**). Furthermore, the cDNA read coverages of the 5′ and 3′ UTRs of *rpsO* and *icd* were higher in the RNA-seq library than in the Ribo-seq library (**Fig. 1C**), showing successful digestion of non-translated or unprotected mRNA regions by RNase I. Similarly, the protein-coding polycistronic *fixN1OQP* mRNA showed high read coverage in the Ribo-seq library along its four ORFs. In contrast, the 5′ leader and 3′ trailer mainly showed coverage in the RNA-seq library, suggesting that they were digested by RNase I (**Fig. 1E**).

**Figure 1.**
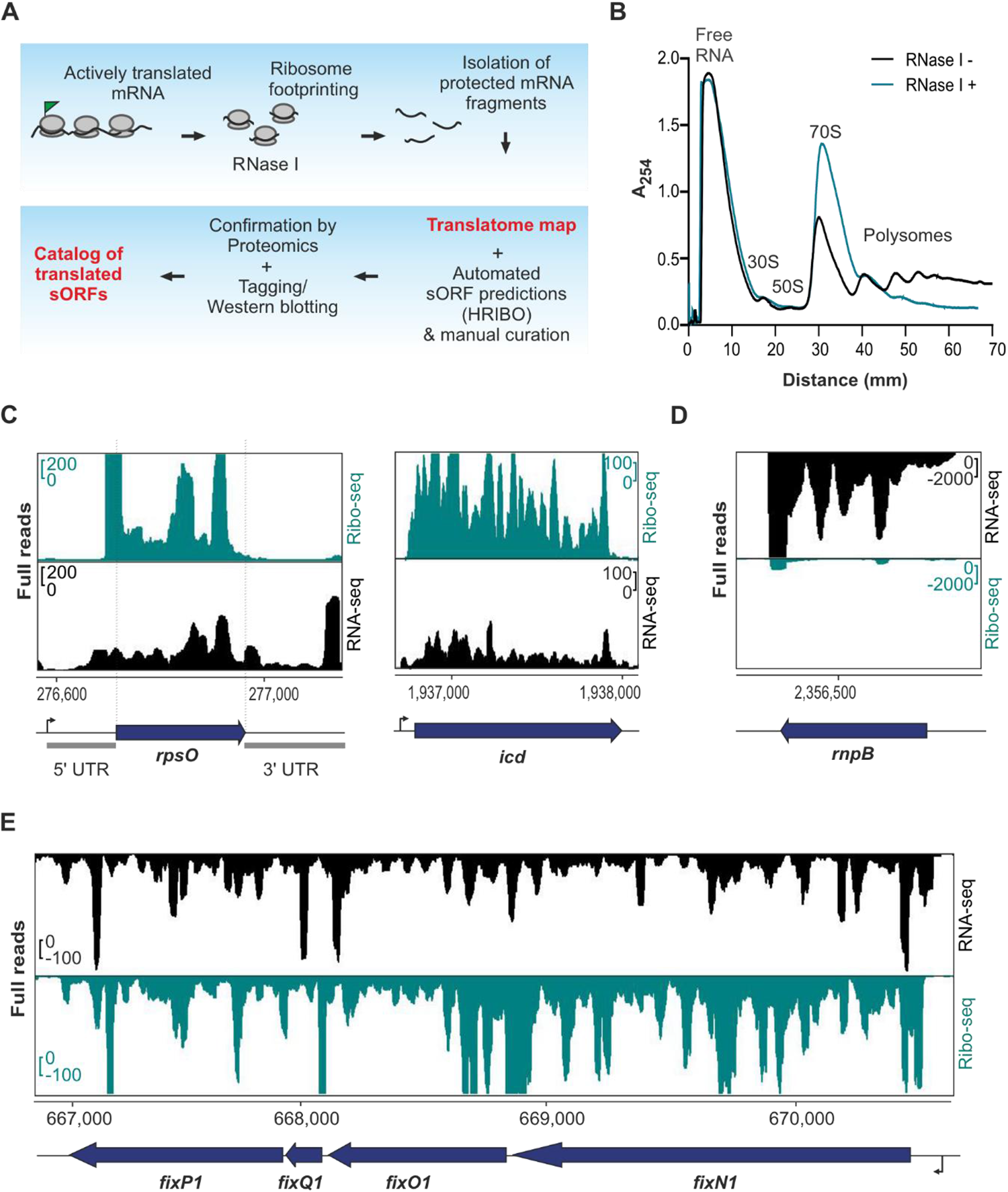
Establishment of ribosome profiling (Ribo-seq) for *Sinorhizobium meliloti*. **(A)** Schematic Ribo-seq workflow to map the *S. meliloti* 2011 translatome. Translating ribosomes (indicated by the polysome fraction) were first captured on the mRNAs. Unprotected mRNA regions were digested by RNase I, converting polysomes to monosomes. Approximately 30-nt-long footprints protected by and co-purified with 70S ribosomes were then subjected to cDNA library preparation and deep sequencing to identify the translatome under the used conditions. The small proteome was identified using HRIBO automated predictions and manual curation. Mass spectrometry and Western blot analysis of recombinant, tagged small open reading frame (sORF)-encoded proteins were used to validate the translated sORFs. **(B)** Sucrose gradient fractionation of the lysates. Cells were harvested at the exponential growth phase by a fast-chilling method to avoid polysome run-off. RNase I digestion led to enrichment of monosomes (70S peak in the green profile) in contrast to the untreated sample (Mock, black profile). Absorbance at 254 nm was measured. **(C)** Integrated genome browser screenshots depicting reads from Ribo-seq and RNA-seq libraries for two annotated ORFs: *rpsO* encoding ribosomal protein S15 and *icd* encoding isocitrate dehydrogenase. They show read coverage enrichment in the Ribo-seq library along their coding parts in contrast to the RNA-seq library but not in the ribosome-non-protected regions (UTRs). The UTRs of *rpsO* are marked. **(D)** Read coverage for *rnpB* corresponding to the housekeeping RNase P RNA. Reads are mostly restricted to the RNA-seq library, suggesting that this RNA is not translated. **(E)** The *fixN1OQP* operon shows read coverage in both the RNA-seq library and Ribo-seq library, the latter indicating that this operon contains translated genes. Genomic locations and coding regions are indicated below the image. Bent arrow indicates the transcription start site based on (Sallet et al. 2013).

The high ribosome density in the Ribo-seq library, which covers the 14 and 12-nt-long intergenic regions between *fixN1 – fixO1* and *fixO1 – fixQ1,* probably represents the footprints of ribosomes that terminate the translation of the upstream ORF and initiate the translation of the downstream ORF. Such events are slower than elongation at most codons in an ORF (Oh et al. 2011). The latter example indicates the translation of the sORF *fixQ1*, which encodes a 50 aa protein (**Fig. 1E**).

Metagene analysis of ribosome occupancy near all annotated start codons (i.e., ATG, GTG, and TTG) showed an enriched ribosome density at the −16 nt upstream (mapping of the 5′ ends of the footprints) and at +16 nt downstream (mapping of the 3′ ends of the footprints) (**Fig. S2A and S2B;** note: +1 is the first nucleotide of the start codon), in line with the expected position of initiating ribosomes waiting to engage in elongation. This feature is a characteristic of translated bacterial ORFs identified by Ribo-seq (Mohammad et al. 2019; Oh et al. 2011). In contrast to MNase-generated Ribo-seq libraries in *E. coli* (Mohammad et al. 2019), no differences in the assignment of ribosome position using the 5′ end or 3′ end mapping approaches were observed (**Fig. S2A** and **S2B**). In the Ribo-seq libraries, we consistently recovered footprints between 27–33 nt (mean at 30 nt), with enrichment of ribosome density strongest at the start codon for the 32 nt footprints (**Fig. S2C** and **S2D**).

### Ribo-seq captures the translatome of *S. meliloti* and reveals features at the single gene level

By comparing the signals of the Ribo-seq and RNA-seq libraries, the TE (Ribo-seq/total RNA coverage) can be estimated at a given locus. This method allowed us to derive a genome-wide estimate of the translatome in minimal medium, where 3,758 of the 6,263 annotated coding sequences (CDS) (60%; GenBank Annotation 2014) had a Ribo-seq signal above the arbitrarily chosen TE cut-off of ≥ 0.5 and RNA-seq and Ribo-seq RPKM of ≥ 10 (see **Methods,** **Fig. 2A, Table S6**). In contrast, the ORF prediction tools implemented in HRIBO (Gelhausen et al. 2021, 2022) detected translation for 2,136 of the 3,758 ORFs (57%), suggesting an average performance in predicting long translated ORFs in *S. meliloti* (**Fig. 2A, Table S6**).

Inspection of the TE for different annotated gene classes and untranslated mRNA regions (all CDS, 5′ and 3′ UTRs, non-coding RNAs, and sORFs) revealed that annotated ORFs exhibited a higher mean TE (TE ≥ 1) compared with non-coding genes, such as housekeeping RNA genes (hkRNA, e.g., tmRNA, 6S, ffs, *rnpB* and *inc*A1/2 RNA, mean TE <1) (**Fig. 2B**), again corroborating the ability of our Ribo-seq data to differentiate between coding and non-coding genes. The 5′ UTR regions of translated mRNAs generally had a mean TE of ≥ 1, which possibly resulted from protection from RNase I trimming of the −16 nt region upstream of the start codon by the initiating ribosomes (**Fig. S2**). This feature was particularly prominent in the leader regions of mRNAs with short 5′ UTRs, indicating that they are partially protected from digestion by initiating ribosomes (**Fig. S3A**). In addition, some 5′ UTRs might contain translated upstream sORFs, such as *trpL* upstream of *trpE* (marked in red in **Fig. 2D**) (Melior et al. 2020). Although less pronounced than at the start codon, the translation-terminating ribosome also protects a certain 3′ UTR region from RNase digestion (Oh et al. 2011), explaining the slightly higher mean TE of 3′ UTRs (**Fig. 2B**). Furthermore, a few of the 3′ UTRs might also contain translated downstream sORFs (**Fig. S3B;** Dodbele and Wilusz 2020; Wu et al. 2020), which may explain the slightly higher mean TE of 3′ UTRs.

**Figure 2.**
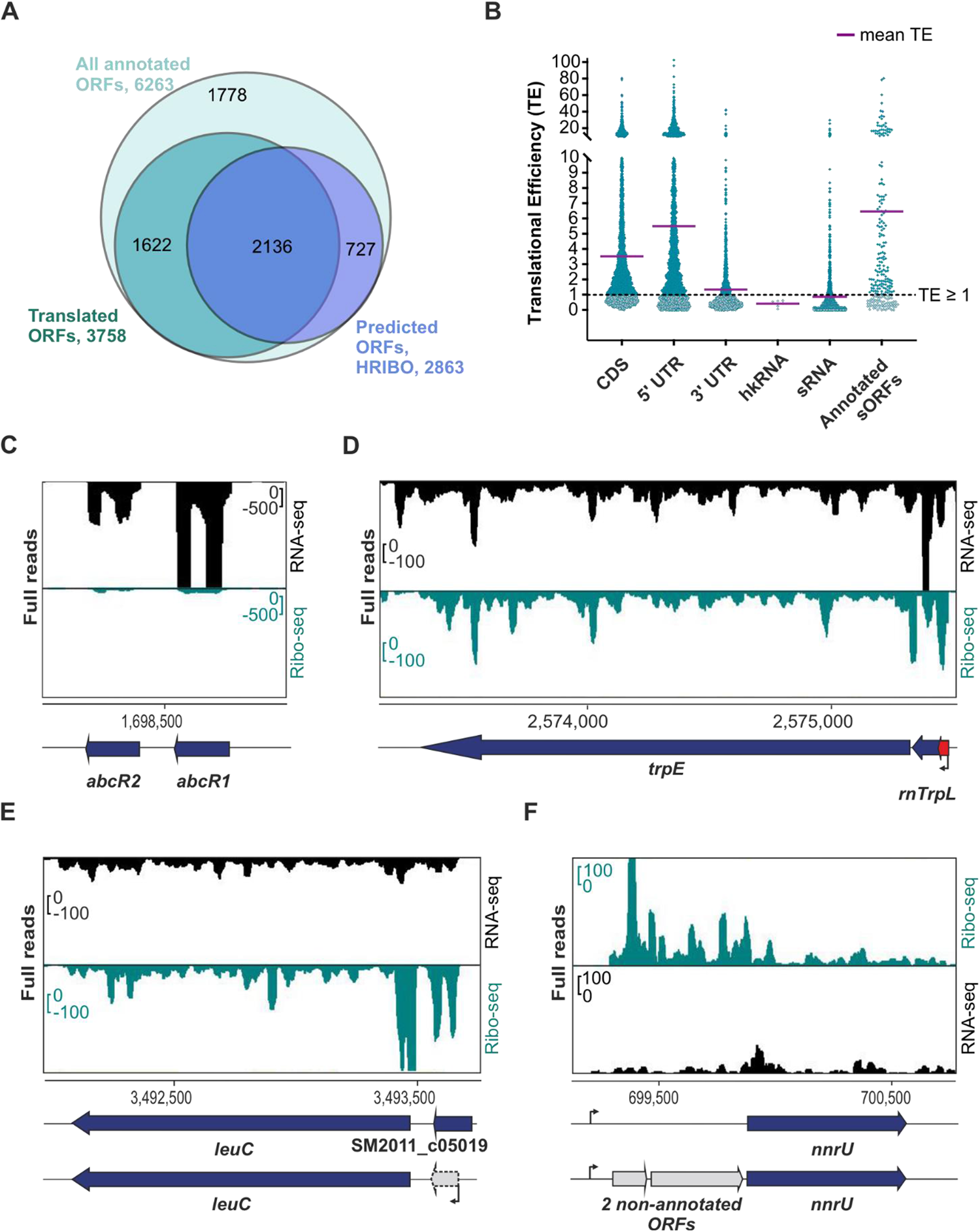
Ribosome profiling (Ribo-seq) captures the translatome of *Sinorhizobium meliloti* 2011 and reveals some features at the single-gene level. **(A)** Comparison of all annotated open reading frames (ORFs), annotated translated ORFs detected by Ribo-seq, and ORFs predicted to be translated by tools included in the HRIBO pipeline. To detect translation, we used the following parameters on the Ribo-seq data: TE of ≥ 0.5 and RNA-seq and Ribo-seq RPKM of ≥ 10. The numbers of ORFs per category are shown and represented by area size. Diagrams were prepared with BioVenn (www.biovenn.nl). **(B)** Scatter plot showing global TEs (TE = Ribo-seq/RNA-seq) computed from *S. meliloti* Ribo-seq replicates for all annotated coding sequences (CDS), annotated 5′ and 3′ UTRs, annotated housekeeping RNAs (hkRNA), annotated small RNAs (sRNAs) with (putative) regulatory functions, and annotated sORFs encoding proteins of ≤ 70 amino acids (aa). The purple lines indicate the mean TE for each transcript class. **(C)** Analysis of the two well-characterized sRNAs AbcR1 and AbcR2 by Ribo-seq. These two sRNAs show read coverage, mostly in the RNA-seq library. **(D)** Ribo-seq reveals the active translation of the *trpE* leader peptide peTrpL (14 aa, encoded by the leaderless sORF *trpL* in the 5′ UTR (red arrow) and/or by the attenuator sRNA rnTrpL). In addition, the coverage of the Ribo-seq library shows that the biosynthetic gene *trpE* is translated in minimal medium, as expected. **(E)** Re-annotation of the annotated sORF SM2011_c05019 (50 aa). The actual annotation does not fit the RNA-seq and Ribo-seq read coverages. HRIBO predicts a shorter leaderless sORF (38 aa) that corresponds to the read coverage in both libraries. **(F)** Two ORFs missing from the GenBank (2014) annotation are revealed by Ribo-seq upstream of the *nnrU* gene related to denitrification. Genomic locations and coding regions are indicated below the image. Bent arrows indicate transcription start sites based on (Sallet et al. 2013).

Most of the annotated sRNAs had a mean TE of < 1, indicating that they are in fact non-coding, such as the sRNAs AbcR1 (TE=0.2) and AbcR2 (TE=0.09) (**Fig. 2C**) (Torres-Quesada et al. 2013). However, some annotated sRNAs had a mean TE of ≥ 1, suggesting that they may be small mRNAs or dual-function sRNAs (**Fig. S3C**). For example, **Fig. 2D** shows the recently described dual-function sRNA rnTrpL (TE = 1.16), which corresponds to the tryptophan attenuator and contains the *trpL* sORF encoding the functional 14 aa leader peptide peTrpL (Melior et al. 2019; Melior et al. 2021). Since rnTrpL is a small, leaderless mRNA starting with the AUG of *trpL*, **Fig. 2D** also exemplifies how our Ribo-seq analysis can capture leaderless translated ORFs. Furthermore, as expected, we detected active translation of the biosynthetic genes *trpE* and *leuC* under growth in minimal medium lacking tryptophan and leucine (**Fig. 2D** and **2E**).

Finally, we used our Ribo-seq data to curate the annotation of *S. meliloti*. For example, Ribo-seq, RNA-seq data, and our computational ORF predictions based on Ribo-seq all indicated that the start of the sORF SM2011_c05019 (50 aa) is likely located downstream of the current annotation, implying a shorter sORF of 38 aa (**Fig. 2E**). Additional sORFs whose annotation should be adjusted are reported in **Table S7**. Moreover, our data revealed additional ORFs that should be added to the genome annotation. For example, the RNA-seq and Ribo-seq read coverages indicate active expression (transcription and translation) upstream of the *nnrU* gene. However, no gene was predicted in this region of the GenBank 2014 annotation. HRIBO’s prediction tools indicated the potential for two non-annotated ORFs encoding 51 and 132 aa proteins upstream of the *nnrU* gene (**Fig. 2F**). The 51 aa ORF is annotated in the related *Sinorhizobium medicae* and *Ensifer adhaeren*s, and in the latter, a homologous 142 aa ORF is annotated between the 51 aa sORF and *nnrU*. Notably, while both ORFs were contained in the *S. meliloti* RefSeq 2017 annotation, the 132 aa ORF was removed again from the latest version (June 2022). This observation underlines the need for and value of integrative approaches that can capture and consolidate reference genome annotations from different annotation centers and even from different releases, which can differ substantially. The iPtgxDB approach (Omasits et al. 2017) represents one strategy to readily capture and visualize such differences, as we show here and for a number of additional cases below.

### Ribo-seq reveals translated annotated small proteins in *S. meliloti*

Among the 6,263 annotated CDS in the *S. meliloti* 2011 genome (the annotation from GenBank 2014 has been used in the laboratory as a reference point for several years), 259 (roughly 4%) correspond to SEPs, with sizes ranging between 30 (the smallest annotated SEP) and 70 aa (**Table S6**). To benchmark our Ribo-seq data for its capacity for global identification of translated sORFs, we analyzed the Ribo-seq read coverage of these 259 annotated sORFs. By applying the TE of ≥ 0.5 and RNA-seq and Ribo-seq RPKM of ≥ 10 cut-off criteria, 131 of them were suggested to be translated (**Table S6**). However, we further included an extensive manual inspection (see **Methods**) of the Ribo-seq read coverage on top of these cut-offs to derive a very high-confidence dataset of 85 (33%) translated sORFs (**Fig. 3A, Table S6**).

We then used this set of manually curated, translated sORFs as a benchmark sORF data set to evaluate the performance of two machine learning-based, automated, Ribo-seq-based ORF prediction tools included in our HRIBO pipeline (Gelhausen et al. 2021, 2022), REPARATION (Ndah et al. 2017), and DeepRibo (Clauwaert et al. 2019). REPARATION predicted the translation for 23 of the 85 benchmark sORFs (26%; **Fig. 3A**), even missing some highly translated sORFs, such as those encoding ribosomal proteins (SM2011_c04434 encoding 50S ribosomal protein L34, mean TE = 5.47) and proteins with housekeeping functions (SM2011_c04884 encoding the anti-sigma factor, mean TE = 2.02, and SM2011_c03850 encoding the heme exporter D, a cytochrome C-type biogenesis protein, mean TE = 0.88). In contrast, DeepRibo predicted translation for 66 of the 85 benchmark sORFs (78%; **Fig. 3A**), indicating that it performed better in terms of detecting translated sORFs from *S. meliloti* Ribo-seq data.

**Figure 3.**
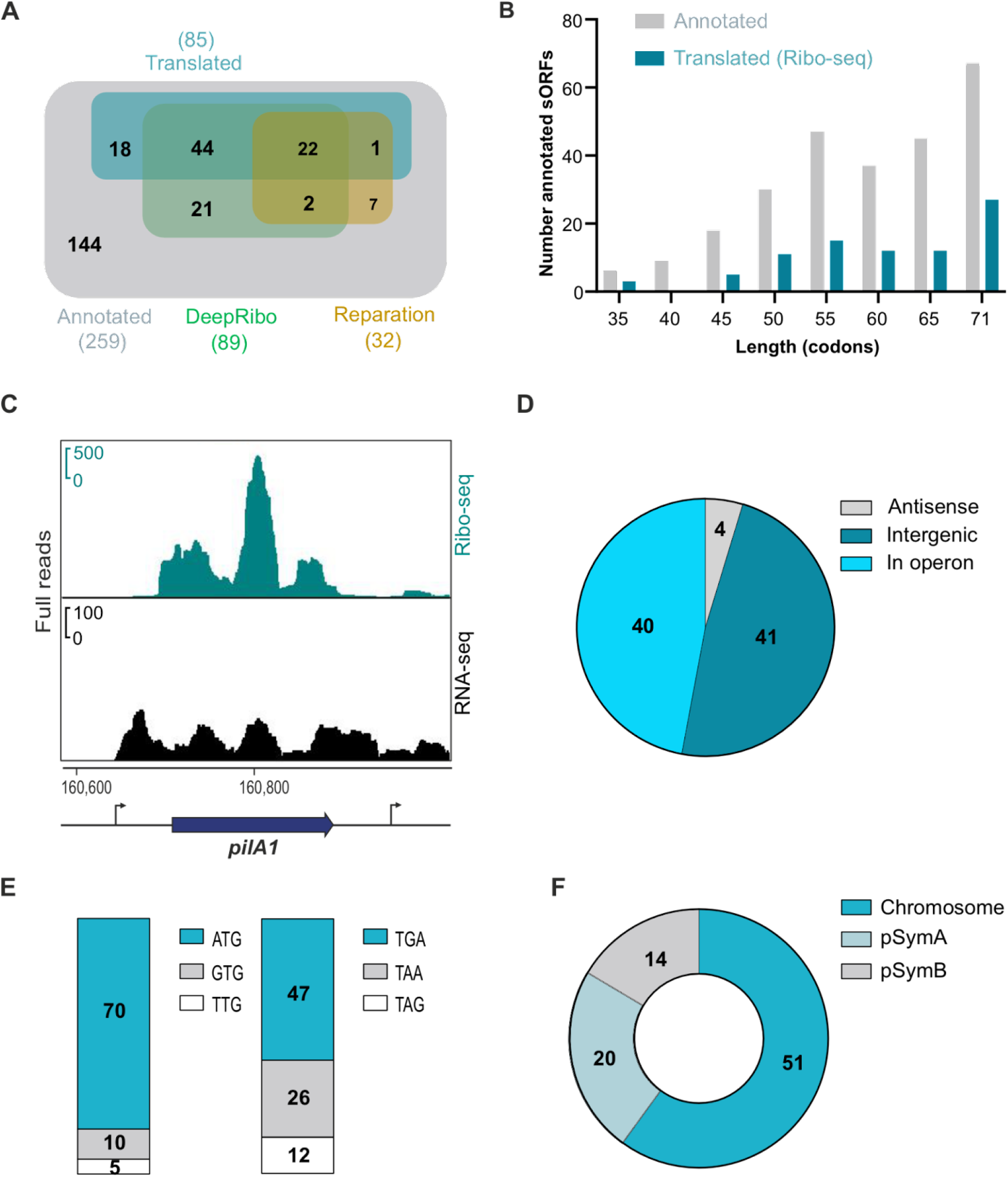
Ribo-seq reveals translated annotated small open reading frames (sORFs) in *Sinorhizobium meliloti* 2011. **(A)** Venn diagrams showing the overlap between all annotated sORFs (259 sORFs, GenBank 2014), the sORFs detected as translated by Ribo-seq (benchmark set, TE of ≥ 0.5, RNA-seq and Ribo-seq RPKM of ≥ 10, and extensive manual curation), and sORFs predicted by the automated ORF prediction tools Reparation or DeepRibo. **(B)** Histogram showing the length distribution of the 85 annotated sORFs identified as translated by Ribo-seq in comparison with the 259 annotated sORFs. **(C)** Integrated genome browser screenshot depicting reads from the Ribo-seq and RNA-seq libraries for the annotated sORF *pilA1* (60 amino acids, encoding a pilin subunit). The genomic position and the coding region are indicated below the image. Bent arrows indicate transcription start sites based on (Sallet et al. 2013). **(D)** Genomic context for the translated annotated sORFs relative to the annotated neighboring genes. **(E)** Start (left) and stop (right) codon usage of the translated annotated sORFs. **(F)** Replicon distribution of the translated annotated sORFs.

The majority of the 259 annotated (76%) and the subset of 85 translated sORFs (78%) encode SEPs of ≥ 50 aa (**Fig. 3B**), in line with the expected poor annotation of very short ORFs.

**Fig. 3C** shows read coverage from the Ribo-seq and RNA-seq libraries for the sORF encoding a 60 aa pilin subunit (TE=23.3), which illustrates the successful digestion of parts of the 5′ and 3′ UTR regions not covered by ribosomes, thus allowing us to define sORF borders. In terms of type of genomic location, most of the translated annotated sORFs are located in intergenic regions and operons, and only a few were found in antisense transcripts (**Fig. 3D**). The vast majority of the translated annotated sORFs were found to start with ATG, followed by GTG and TTG. The stop codon preference, although less pronounced, was TGA > TAA > TAG (**Fig. 3E**). Finally, 60% of the 85 translated annotated sORFs were located on the chromosome, 23.5% on the megaplasmid pSymA, and 16.5% on the megaplasmid pSymB (**Fig. 3F**).

### Ribo-seq further expands the small proteome of *S. meliloti*

We then aimed to exploit the sensitivity of Ribo-seq to identify potential novel *S. meliloti* 2011 sORFs missing from GenBank annotation (2014) and thereby provide a nearly complete catalog of its small proteome. The two machine learning-based, automated, Ribo-seq-based ORF prediction tools integrated into the HRIBO pipeline produced a large number of predictions (approximately 15,000) for potential non-annotated sORFs (**Fig. 4A**), as previously shown in other bacterial species (Gelhausen et al. 2022). Given that these ORF prediction tools neither consider RNA-seq data nor TE but only utilize ribosome occupancy, we decided to filter the predictions for those with RNA-seq and Ribo-seq RPKM values of ≥10 and mean TE of ≥ 0.5. In addition, we applied a stringent cut-off for the DeepRibo score (see **Methods**) that allowed the ORF candidate ranking, which led to 266 candidates of translated non-annotated sORFs. Manual curation of all candidates based on their Ribo-seq coverage left us with a list of 54 non-annotated sORFs, which we proposed with high confidence to be translated during growth of *S. meliloti* in minimal medium (**Fig. 4A; Table S7**). Overall, the 54 non-annotated sORFs were shorter than the annotated ones: 33 of them (61%) correspond to SEPs with lengths between 10 and 49 aa, and nine of them (17%) represent SEPs shorter than 30 aa (the shortest annotated ORF in the *S. meliloti* annotation). A comparison of the length distribution of the 85 annotated and 54 non-annotated translated sORFs (**Fig. 4B**) illustrates the potential of Ribo-seq to detect very short translated sORFs.

The 54 non-annotated sORFs are encoded in diverse genomic contexts (**Fig. 4C**): 33% were located on annotated sRNAs, suggesting that these sRNAs are small mRNAs or dual-function sRNAs, 26% were in the intergenic regions, thus defining small mRNAs, and 20% were in the 5′ UTRs and may correspond to regulatory upstream ORFs (Evguenieva-Hackenberg 2022). Only a few were located in 3′ UTRs, on antisense transcripts and in an operon (**Fig. 4C**). Moreover, the majority of the 54 sORFs (63%) were located in the chromosome, 22% on pSymA, and 15% on pSymB (**Fig. 4D**), a distribution comparable to that of the annotated sORFs (**Fig. 3F**). Similar to the annotated sORFs, ATG was also the preferred start codon among the 54 non-annotated translated sORFs, and only five and four sORFs started with GTG or TTG, respectively; their stop codon preference was also similar to that of the annotated sORFs (**Table S6**). Importantly, as the iPtgxDB integrates and consolidates different reference genome annotations and various predictions, we could readily deduce that 11 of the 54 translated sORFs were contained in the 2017 RefSeq annotation, precisely matching their predicted start and stop codons (**Table S7**). Five candidates matched a RefSeq annotation, but they were shorter. One candidate matched the stop but was only 1 aa longer than the RefSeq annotation. Finally, three candidates matched a GenBank stop codon, but they were shorter than annotated (one of which was in fact again removed in the RefSeq annotation). In summary, Ribo-seq uncovered 37 translated sORF candidates that were novel compared to both in GenBank 2014 and RefSeq 2017 annotations (**Table S7**).

**Figure 4.**
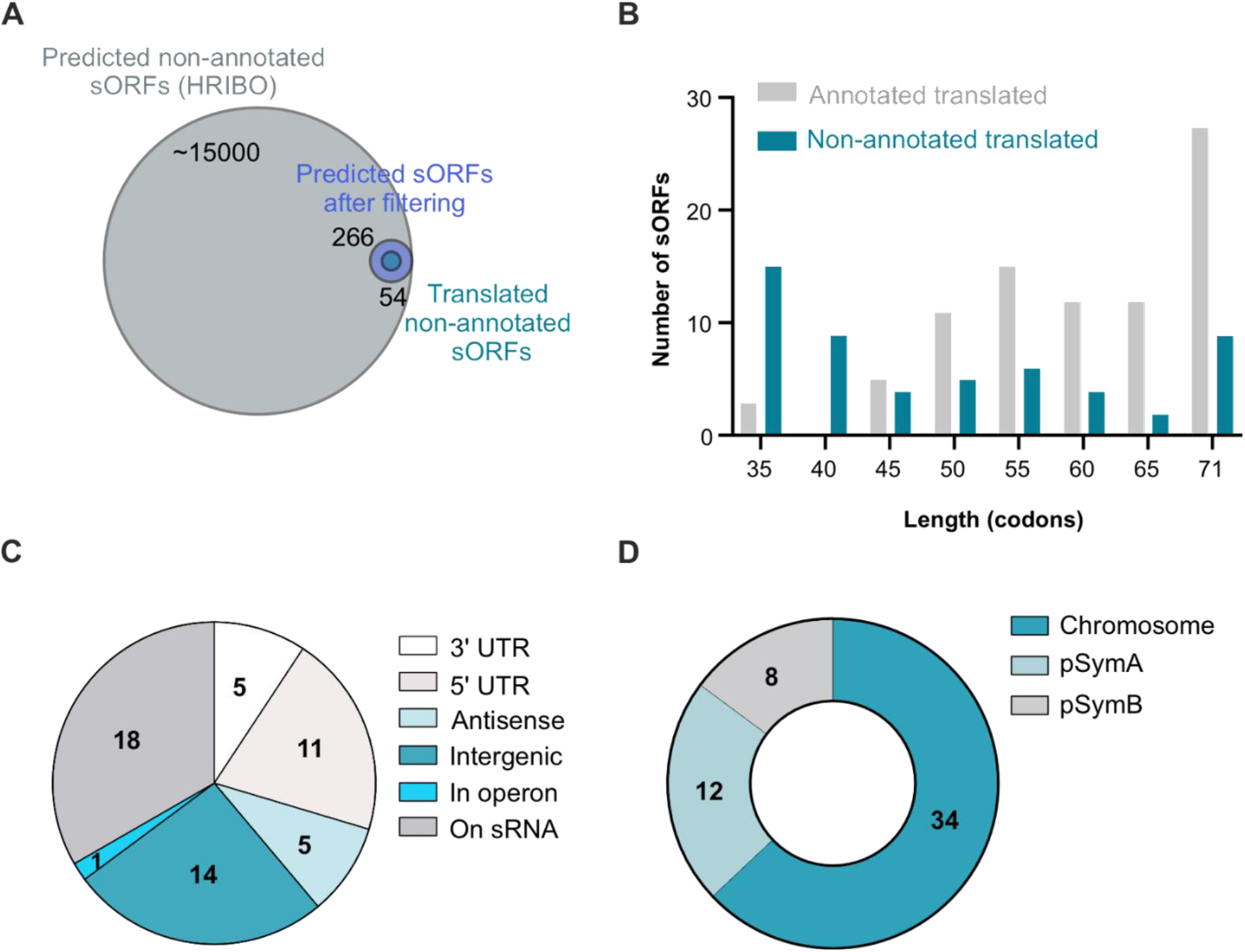
Ribo-seq uncovers a repertoire of small open reading frames (sORFs) missing from the *Sinorhizobium meliloti* 2011 genome annotation. **(A)** sORF predictions from HRIBO included a high number of potential non-annotated sORFs (approximately 15,000). These sORFs were first filtered (TE of ≥ 0.5, RNA-seq and Ribo-seq RPKM of ≥ 10, DeepRibo score of > −0.5) to generate a set of 266 translated sORF candidates that were additionally manually curated by inspection of the Ribo-seq read coverage in a genome browser. Overall, 54 high-confidence non-annotated sORFs displayed active translation during growth in minimal medium. A Venn diagram shows the respective number of proteins from each category (scaled with area size). Diagrams were prepared with BioVenn (www.biovenn.nl). **(B)** Histogram showing the length distribution of the 54 non-annotated versus the 85 annotated sORFs identified as translated by Ribo-seq. **(C)** Genomic context of the translated non-annotated sORFs. **(D)** Replicon distribution of the translated non-annotated sORFs.

### Both standard and small custom iPtgxDBs informed by Ribo-seq data facilitate novel SEP identification by MS

To validate sORF translation and identify novel SEPs of *S. meliloti* 2011, we then conducted MS-based proteomics using experimental strategies to increase the coverage of the MS-detectable small proteome and two search DBs. Cells were cultured either in minimal GMS medium (same as for Ribo-seq) or in rich TY medium, and three complementary sample preparation approaches were used: 1) tryptic in-solution digest of all proteins (a standard proteomics approach), 2) solid phase enrichment (SPE) of small proteins with subsequent Lys-C digestion, and 3) SPE of small proteins without subsequent digestion (**Fig. 5A**). Approaches 2 and 3 can identify SEPs whose peptides are not within the detectable range (approximately 7 aa to 40 aa) upon a tryptic digest (Tyanova et al. 2016).

For the DB searches, we first relied on a standard (full) iPtgxDB (Omasits et al. 2017) that hierarchically integrates reference genome annotations, *ab initio* gene predictions, and *in silico* ORF predictions (see **Methods**). The overlap and differences of all annotation sources were captured and consolidated in a composite gene identifier. Moreover, a large but minimally redundant protein search DB (for more details, see https://iptgxdb.expasy.org/creating_iptgxdbs/) is created, as well as a GFF that allows researchers to overlay experimental evidence, such as RNA-seq, Ribo-seq, or proteomics data. Individual iPtgxDBs must be prepared for different proteases (see **Methods**). For trypsin, the standard iPtgxDB contained close to 160k protein entries of approximately 103k annotation clusters (**Table S3.1**), that is, genomic loci that share the stop codon but have different predicted protein start sites. Approximately 92% of the peptides unambiguously identify one protein entry, which are called class 1a peptides that facilitate downstream data analysis and allow to swiftly identify novel proteoforms or SEPs. Although standard iPtgxDBs are very large, when combined with stringent FDR filtering, they have provided convincing results in the past, that is, for the identification of novel SEPs that withstood independent validation efforts (Omasits et al. 2017; Melior et al. 2020; Bartel et al. 2020). However, as large DBs inflate the search space, they complicate protein inference and FDR estimation, resulting in a large likelihood of a random hit, especially for SEPs (Fancello and Burger 2022; Nesvizhskii 2010). Importantly, the 266 top Ribo-seq-implied novel candidates (**Fig. 4A**) allowed us to explore whether a much smaller custom iPtgxDB may provide additional value for the identification of annotated or novel SEPs. Adding these 266 candidates to the three reference genome annotations (RefSeq, GenBank, Genoscope) and the Prodigal *ab initio* gene predictions resulted in a 20-fold smaller custom iPtgxDB (**Fig. 5A**) (approximately 8,000 protein entries in 7,300 annotation clusters), with a higher percentage of class 1a peptides (nearly 98%; **Table S3.3**).

**Figure 5.**
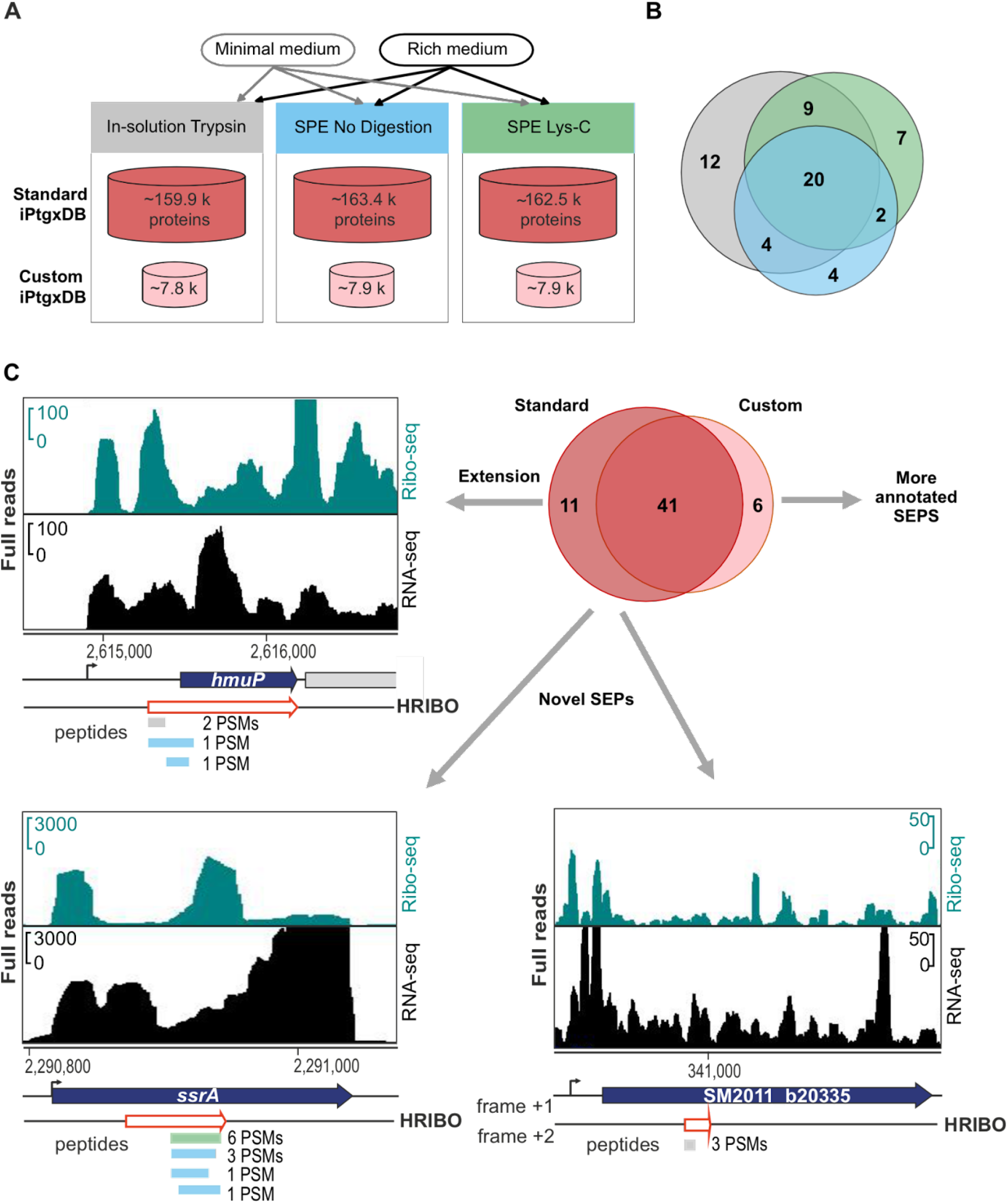
Mass spectrometry-based identification of known and novel small open reading frame-encoded proteins (SEPs). **(A)** Experimental set-up for the proteomics analyses. Bacteria were grown in minimal and rich media, and protein extracts were further processed with tryptic in-solution digest (gray), solid-phase enrichment (SPE) of small proteins with subsequent Lys-C digestion (green), or without further digestion (blue). **(B)** Overlap of the identified SEPs by experimental approach; trypsin identified 45 SEPs; compared with the trypsin approach, Lys-C identified 38 SEPs (nine novel, 24%), and the approach without digestion found 30 SEPs (six novel, 20%). **(C)** Novel/unique identifications uncovered by the standard integrated proteogenomic search databases (iPtgxDB) and the small custom iPtgxDB. Standard iPtgxDB: Three peptides imply a 14 aa longer proteoform (60 aa) for HmuP than annotated; four peptides of the tmRNA-encoded proteolysis tag were identified; one peptide (3 peptide spectrum matches [PSMs]) implied a novel SEP (34 aa) internal to the genomic region that also encodes SM2011_b20335 but in a different frame. Spectra identifying these peptides are shown in **Fig. S5**. These identifications were also predicted by HRIBO based on Ribo-seq. Finally, six annotated proteins (GenBank 2014 and/or RefSeq 2017) were identified only in the search against the small custom iPtgxDB, as they did not accumulate enough spectral evidence in the search against the standard iPtgxDB (**Table S4**).

The acquired MS-spectra were searched against the standard and small iPtgxDBs, and the results were compiled and stringently filtered, requiring more PSM evidence (see Methods) for *ab initio* and *in silico* predictions (Varadarajan et al. 2020a; Varadarajan et al. 2020b). Overall, more than 1,200 annotated proteins were detected at an estimated protein FDR of approximately 1%. The SPE-based small protein enrichment steps uniquely identified 160 of these proteins (**Fig. S4A**). Notably, the search against the small custom DB accounted for 112 unique identifications (**Fig. S4B**), presumably due to improved search statistics. The MS-identified proteins included 58 SEPs, with ≤ 70 aa, 47 of which were annotated (GenBank 2014 and/or Refseq 2017) (**Table S4**). Similar to the overall results, the two SPE approaches also added unique SEPs: while 45 of the 58 MS-detected SEPs were identified with standard trypsin-based digestion, 13 SEPs were uniquely identified after processing the samples with SPE and either a Lys-C digest (9 of 38 not covered by trypsin) or no proteolytic digest (6 of 30 not covered by trypsin) (**Fig. 5B**). Most MS-identified SEPs were between 60 and 70 aa long (67%), and the smallest detected SEP was 20 aa long. They include abundantly expressed proteins (the cold shock proteins RS25125 and RS00515, and RS31025, a 50S ribosomal protein L32) (**Table S4**) down to candidates identified by only 2 PSMs, such as a 59 aa hypothetical protein, which we refer to as SEP7 (see next section). Among the 85 GenBank-annotated SEPs identified with high confidence, as translated by Ribo-seq (**Fig. 3A**), 31 were identified by MS. Among the 54 SEPs missing from the GenBank annotation and identified as translated by Ribo-seq (**Fig. 4A**), five were identified by MS, and those are present in the Refseq 2017 annotation (**Table S4**).

Importantly, both searches added unique identifications. The search versus the full iPtgxDB added 11 potential novel SEPs or longer proteoforms than annotated, which were *in silico* predictions that were excluded from the small custom iPtgxDB. A 14 aa longer proteoform of HmuP was identified by three peptides with 4 PSMs (**Fig. S5A**). Here, when manually inspecting the Ribo-seq data, it perfectly agreed with the extension of the 46 aa GenBank annotation (**Fig. 5C**). This finding exemplifies how proteomics and Ribo-seq jointly identify a novel proteoform. Furthermore, the tmRNA-encoded 12 aa proteolysis tag peptide was uniquely identified, which marks incompletely translated proteins for degradation (Karzai et al. 2000) (**Fig. 5C**). The tag peptide was identified as a C-terminal part of an *in silico* predicted 23 aa SEP included in the standard iPtgxDB. It was only detected in the minimal medium by four peptides: one in the Lys-C digest and three from the search without protease (**Fig. 5C** and **S5B**). Mutation of the start codon of the 23 aa sORF had no effect on the translation of the proteolysis tag peptide, in line with the mechanism proposed for this split tmRNA (Ulvé et al. 2007; Keiler et al. 2000) (**Fig. S6**). An example of a completely novel 34 aa SEP is shown in the third panel of **Fig. 5C** (see also **Fig. S5C**); it is located in a genomic region that harbors an annotated CDS and is translated in a different frame. The novel sORF has Ribo-seq support (TE 0.4) but did not pass our stringent Ribo-seq cut-offs. Notably, the search against the small custom iPtgxDB added six unique SEP identifications (again, due to better search statistics) (**Fig. 5C**). Four of them were also among the 85 GenBank-annotated SEPs identified by Ribo-seq data (RS33030, RS33620, RS33980, and a6027), lending independent support for their expression (**Table S4**). RS33620 belongs to the arginine-rich DUF1127 family of proteins, the members of which are involved in phosphate and carbon metabolism in *Agrobacterium tumefaciens* (Kraus et al. 2020), and in RNA maturation and turnover in *Rhodobacter* (Grützner et al. 2021). In addition, the abovementioned RefSeq-annotated SEP7 was identified (**Fig. S5D**). Two other SEPs (one of them novel) were identified with only 1 PSM (**Fig. S5E** and **S5F**), which was below our threshold, but had strong Ribo-seq support (SEP1, SEP20; see next section).

### Validation of a subset of Ribo-seq-implied small proteins by Western blot analysis

Since out of the 54 high-confidence Ribo-seq-implied sORFs that were not contained in the GenBank annotation (2014) only five were detected with at least 2 PSMs in the MS analysis (**Table S4**), we attempted validation by epitope tagging and Western blot analysis (**Fig. 6**). Nineteen sORFs were selected that i) cover a broad range of TE values, ii) start with one of the three main start codons (ATG: 16 sORFs, GTG: two sORFs, or TTG: one sORF), and iii) were either added in the RefSeq 2017 annotation (five sORFs) or were novel with respect to these two annotations (14 sORFs). The corresponding proteins were designated SEP1 to SEP19 (**Table S4**). They included three of the candidates that were also detected by MS (SEP7: 2 PSMs; SEP 10: 59 PSMs; SEP17: 29 PSMs). SEP1 was only identified by 1 PSM, that is, below the threshold (**Table S4; Fig. S5E**), but with strong Ribo-seq support (highest TE among the 54 high-confidence Ribo-seq candidates; **Table S7**). Moreover, three SEP candidates below 30 aa (SEP1, SEP3, and SEP6) and four candidates with a predicted transmembrane helix (TMH) (SEP4, SEP6, SEP13, and SEP16; **Table S4**) were analyzed. As a 20th candidate (SEP20), we included a conserved annotated sORF located in the cytochrome C oxidase cluster *ctaCDBGE* between *ctaB* and *ctaG* (GenBank annotation 2014), which also contains a predicted TMH. SEP20 was identified by 1 PSM in the MS analysis (**Fig. S5F**) and did not pass the stringent HRIBO criteria for translated candidate sORFs (**Table S3,** TE = 6.99, RPKM of < 10 in replicate 1) but showed strong read coverage in the Ribo-seq library (**Table S6**).

Each sORF was cloned together with its −15 nt 5′ UTR region, thus containing its putative ribosome binding site in frame to the SPA-tag encoding sequence into plasmid pSW2 (**Fig. 6A** and **Fig. S1**). Transcription of the sORF::*spa* fusion is under the control of a *S. meliloti sinI* promoter (P*sinI*) of moderate strength, which is constitutively active (Charoenpanich et ^a^l. 2013). Thus, the detection of a SEP-SPA fusion protein by Western blot analysis would indicate sORF translation. The Western blot analysis of crude lysates of cultures grown in minimal medium using FLAG-directed antibodies revealed signals for 15 of the 20 candidates, including SEP20 (see **Fig. 6B–6G**). For the 12 candidates, one band consistent with their predicted SEP length was detected. For SEP1 and SEP5, on top of the expected SEP-SPA bands, slow migrating bands at approximately 25 kDa (see asterisks in **Fig. 6E**) were detected, which probably corresponded to a non-specific signal, as they were also detected in some EVC samples after lysate fractionation (**Fig. S7**).

**Figure 6.**
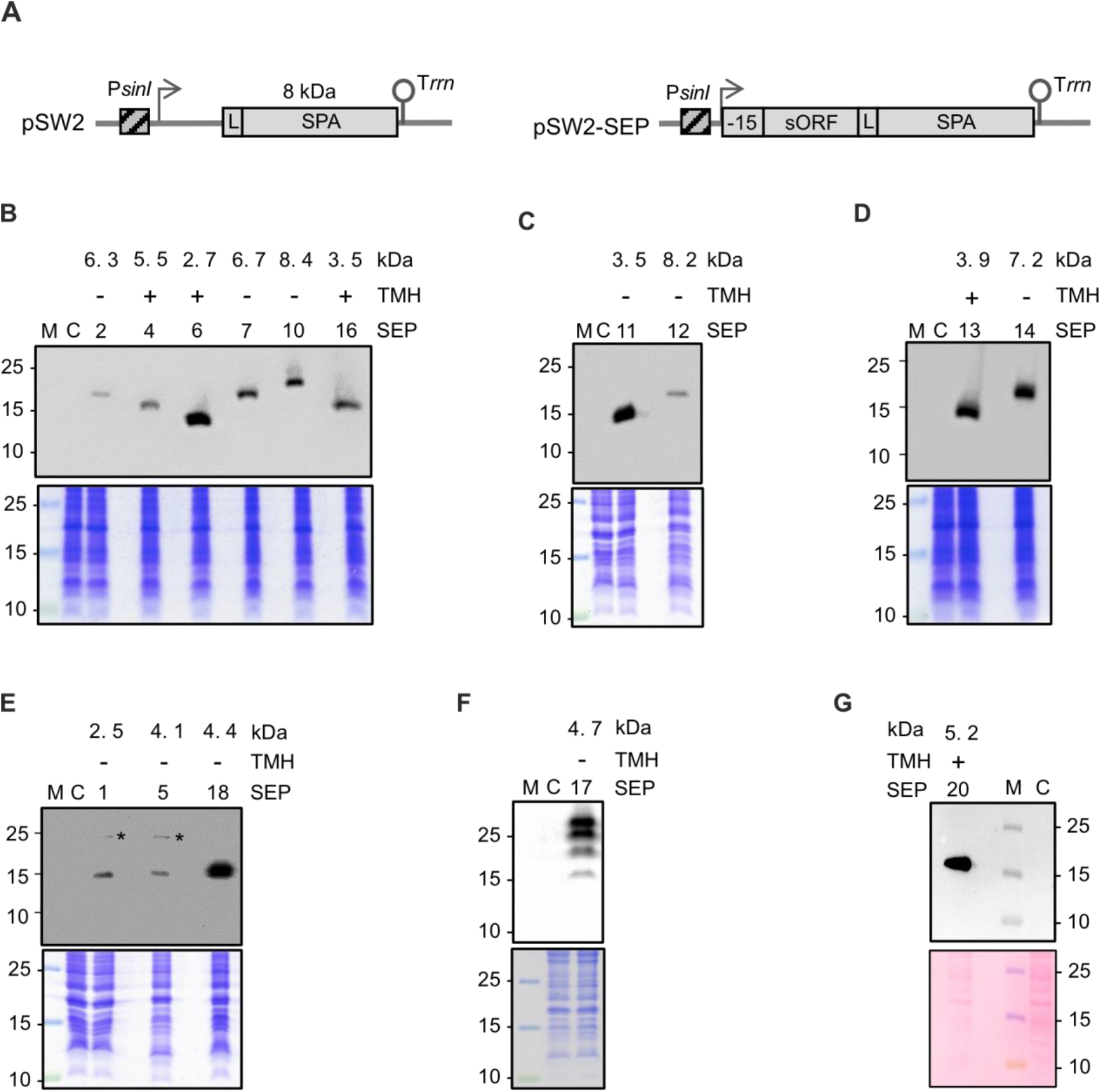
Detection of 15 sequential peptide affinity (SPA)-tagged small open reading frame-encoded proteins (SEPs) in *Sinorhizobium meliloti* crude lysates. **(A)** Schematic representation of the empty plasmid pSW2 (contains no promoter and no ribosome-binding site upstream of the linker [L] and SPA-encoding sequence) and a pSW2-SEP plasmid for the analysis of sORF translation. The constitutive P*sinI* promoter (hatched box), the corresponding TSS (flexed arrow), the sORF coding sequence with its −15-nt-long region, the SPA-tag (with its molecular size indicated) preceded by a linker (L) (gray boxes), and the T*rrn* terminator (hairpin) are depicted. (B) to (F) Western blot analysis of crude lysates (upper panels) and the corresponding Coomassie-stained gels, and (G) corresponding Ponceau-stained membrane for selected SEPs. Monoclonal FLAG-directed antibodies were used. Migration of marker proteins (in kDa) is shown on the left side. * Unspecific signal. Above the panels, the numbers of the analyzed SEP protein (see Table S7), the presence (+) or absence (−) of a predicted TMH, and the molecular size (in kDa) of the SEP without the SPA tag are given. M: protein marker. C: empty vector control, lysate from a strain containing pSW2.

The bands of the tagged SEP1 and SEP5 ran similarly, although SEP1 is smaller than SEP5, as indicated above the panel (**Fig. 6E**). Probably, the aberrant migration of SEP1 is due to its acidic aa composition (pI of 4.18) (Guan et al. 2015). SEP17 showed multiple bands, with a weak and fast migrating band at approximately 15 kDa, which probably corresponds to the monomeric SEP17-SPA protein, and three strong and slow migrating bands, which could indicate protein oligomerization (**Fig. 6F**). Overall, the translation of SEPs with alternative start codons, that is, GTG (SEP10 and SEP14) and TTG (SEP7), and of the five candidates missed in the GenBank (2014) annotation but included by Refseq (2017) (SEPs Nr. 4, 7, 10, 17, and 18), was validated. Importantly, this analysis confirmed the translation of six novel SEPs (SEPs Nr. 1, 6, 11, 13, 14, and 16), including two of the three SEP candidates shorter than 30 aa. Finally, our observation that 11 (out of 16) sORFs without MS support but with high confidence Ribo-seq data were validated in the Western blot analysis shows the power of Ribo-seq to detect novel translated sORFs.

Since the analysis of exclusive or predominant subcellular localization is valuable for linking hypothetical proteins without any annotation to some potential function (Stekhoven et al. 2014), we decided to investigate the subcellular localization of the validated SPA-tagged SEPs by Western blot analysis of the supernatant (S100) and pellet (P100) fractions (see Methods) (**Fig. S7**). As expected, the predicted TMH-containing proteins SEP4, SEP6, SEP13, SEP16, and SEP20 were detected exclusively or predominantly in P100, which contains ribosomes and membranes, whereas the predicted cytoplasmic proteins SEP5 and SEP12 were detected exclusively in the S100 fraction **(Fig. S7**). The remaining eight SEPs were detected exclusively or partially in the P100 fraction, suggesting that they could be associated with membrane complexes or ribosomes (SEP10 and SEP18 show similarities to the ribosomal proteins S21 and L7/12) or be prone to aggregation in their recombinant, tagged form.

### Conservation and potential functions of *S. meliloti* novel small proteins

As described above, we detected the translation of 48 sORFs missing in the GenBank 2014 and Refseq 2017 annotations (37 identified by Ribo-seq and additional 11 by MS), which we refer to as novel. Since conserved SEPs are likely to be functional, we used tBLASTn (Gertz et al. 2006) to examine the conservation of the proteins encoded by the 48 novel sORFs (**Fig. 7; Table S8**). The tBLASTn searches were conducted in bacteria with parameters previously established to identify conserved bacterial sORFs (Allen et al. 2014) (see **Methods**). We found a wide range of conservation, from an sORF detected in only four *S. meliloti* strains overall, to sORFs conserved at different higher taxonomic levels, to highly conserved sORFs present in different bacterial phyla (**Fig. 7**). Among the 14 sORFs encoding SEPs with < 30 aa (excluding the tmRNA sORF64), four are conserved beyond *S. meliloti*. One of the most widely conserved novel SEPs is a 64-aa-long small protein detected only by MS (sORF61 in **Fig. 7**). It was identified as a product of an *in silico* predicted sORF, with 3 PSMs in lysates from MM cultures (**Fig. S5G; Table S4**). However, no expression at the level of RNA was detected at its locus, possibly suggesting high protein stability. sORF61 has homologs in several bacterial phyla and multiple paralogs, with a maximal aa sequence identity of 64% on each replicon in *S. meliloti* 2011. Despite its wide distribution and strong conservation, its function is unknown. Overall, excluding the tmRNA, we detected seven sORFs conserved beyond the family *Rhizobiaceae,* suggesting that the corresponding SEPs may have important general functions.

Furthermore, we used TMHMM (Krogh et al. 2001) and PSORTb (Yu et al. 2010) to predict the presence of transmembrane helices and the subcellular localization of the 48 novel SEPs. Localization in the cytoplasmic membrane was predicted for seven SEPs using at least one of the tools (**Fig. 7; Table S8**). Among them are the Ribo-seq-identified and Western blot analysis-validated SEP6 (prediction by both TMHMM and PSORTb) and SEP16 (prediction by TMHMM only), which were detected with strong signals predominantly in the P100 fraction (see **Fig. S7**). The corresponding sORF6 and sORF16 are conserved in *Hyphomicrobiales* (**Fig. 7**). No proteins with predicted membrane localization were found among the 11 MS-detected SEPs (**Fig. 7**). Notably, two of the 48 novel SEPs harbor a predicted SpII cleavage site and are thus probably lipoproteins (**Fig. 7**). Lipoproteins play important roles in physiology, signaling, cell envelope structure, virulence, and antibiotic resistance (Kovacs-Simon et al. 2011); however, as we had previously reported, they are often missed in prokaryotic genome annotations (Omasits et al. 2017).

**Figure 7.**
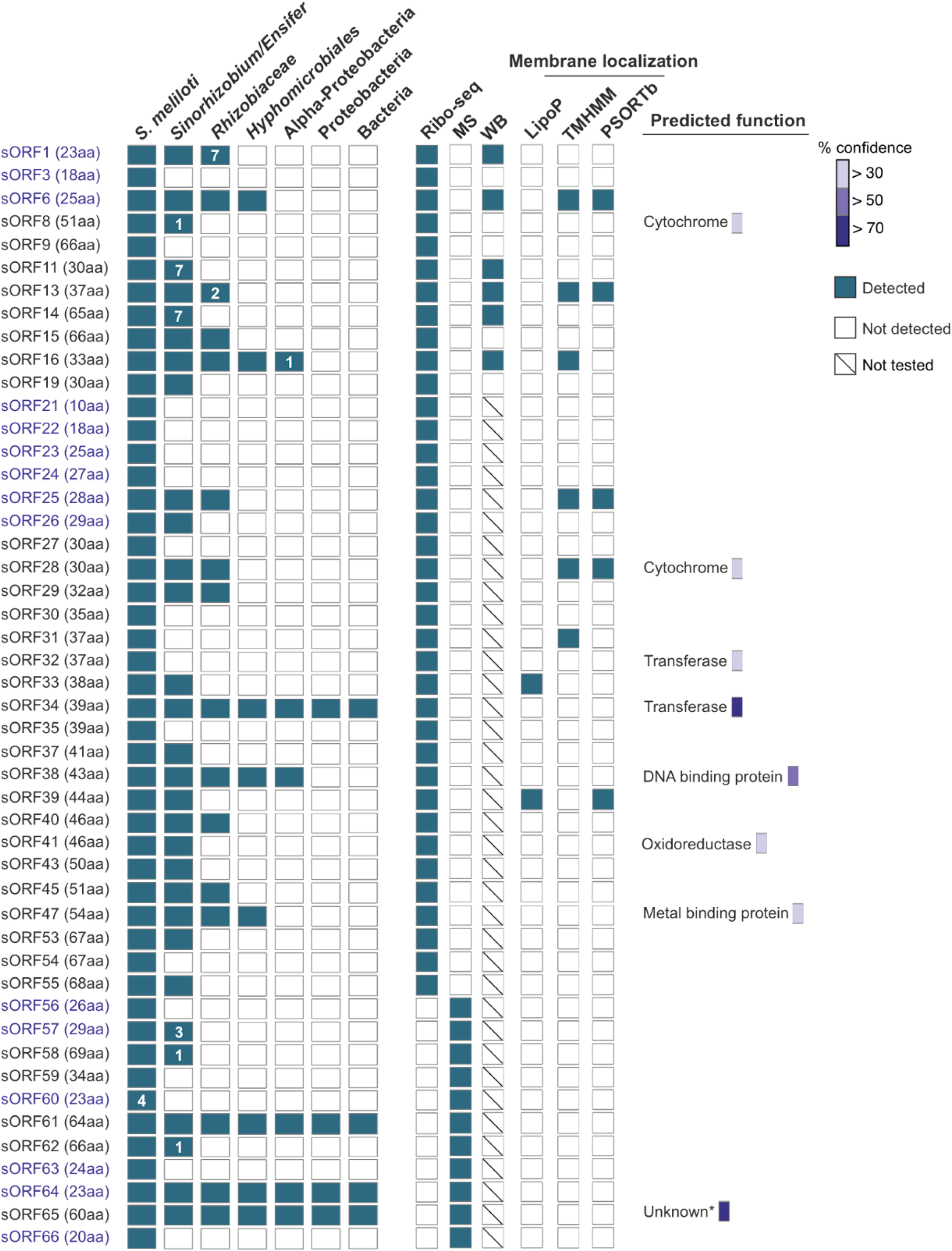
Conservation analysis and functional prediction for 48 novel small open reading frames (sORFs) of *Sinorhizobium meliloti* 2011. The conservation analysis was conducted using tBLASTn. The respective hits (see methods for parameters and cutoffs) are broadly summarized at the level of different taxonomic groups. The number of species outside the lower taxonomic unit, which harbors a hit, is given, if at < 10. In addition, the method by which the respective sORF was detected or confirmed is shown (Ribo-seq: ribosome profiling, MS: proteomics, WB: Western blot), as well as the results of predictions for membrane localization (by TMHMM and PSORTb), signal peptide II cleavage sites of lipoproteins (by LipoP), and function (by Phyre2; only hits with confidence levels greater than 30% are shown). sORF1 to sORF55 are a subset of the 57 Ribo-seq-detected, translated sORFs, which are listed in Table S7, and sORF56 to sORF66 represent the novel sORFs identified by proteomics. sORFs encoding small proteins below 30 amino acids are shown in blue. The putative sORF64, present in tmRNA, contains the proteolytic tag sequence. The sORF65 corresponds to the N-terminal HmuP extension; outside of Proteobacteria, it is conserved in many genera of Planctomycetes. *Structural genomics (92% confidence homology to protein of unknown function).

Moreover, we used Phyre2 (Kelley et al. 2015) to gain insights into the potential functions of novel SEPs with ≥ 30 aa by analyzing their similarity to proteins with known tertiary structures (**Fig. 7; Table S8**). Best hits with a confidence homology of ≥ 30% were obtained for eight novel SEPs (**Fig. 7**). The highest confidence homology suggesting a function was obtained for the SEPs encoded by sORF38 (DNA binding; 18 of the 43 aa residues were modeled with 66% confidence homology; conserved in Alphaproteobacteria) and sORF34 (bleomycin resistance; 37 of the 39 aa residues were modeled with 92% confidence homology; conserved among Bacteria). The HmuP extension (sORF65 in **Fig. 7**; see also **Fig. 5C**) was modeled with 98% confidence along 59 of its 60 aa residues; however, according to Phyre2, the function of the hit was unknown. Overall, obtaining clear functional predictions was not possible even for conserved SEPs, most probably due to their small size.

We also suggest the functions for three annotated sORFs/SEPs with validated translation. SEP5 (added in the RefSeq 2017 annotation) is conserved only in *Sinorhizobium*. Its translation was detected with Ribo-seq and Western blot analysis (**Fig. 6E; Table S4**). The SEP5 sORF contains a cluster of six threonine and three lysine codons near its 3′ ‘end and is located in the 5′ UTR of the aspartate dehydrogenase-encoding gene. Since aspartate is a part of the threonine and lysine biosynthesis pathway (Vitreschak et al. 2004), our observation suggests that this sORF can be involved in the post-transcriptional regulation of the aspartate dehydrogenase gene in *Sinorhizobium*, and SEP5 is possibly a leader peptide. Furthermore, among the annotated SEPs with functional assignment, which were detected by MS, an entericidin A/B family lipoprotein was found (CP004140.1:2141558-2141716:-, **Table S4**). The 52-aa-long protein has a predicted TMH and is conserved in Alphaproteobacteria. Its *A. tumefaciens* homolog, the lipoprotein Atu8019, is involved in specific cell–cell interactions as a part of outer membrane vesicles (Knoke et al. 2020). Finally, an annotated small protein validated in this work by Ribo-seq and Western blot analysis (1 PSM in the MS) is the abovementioned SEP20 (**Fig. 6G, Fig. S5E; Table S4**). It contains a predicted TMH and is conserved in the Alphaproteobacteria, and its sORF is part of an uncharacterized cytochrome oxidase operon. This synteny suggests that SEP20 can participate in the assembly and/or function of the corresponding cytochrome oxidase complex, as previously shown for SEP CydX and cytochrome bd oxidase in *Brucella abortus* (Sun et al. 2012).

## DISCUSSION

In this work, we have developed and applied a Ribo-seq workflow to comprehensively map the translatome of *S. meliloti* 2011 under free-living conditions in a minimal medium. By combining Ribo-seq and MS-based proteomics in a proteogenomic approach, we added 48 novel SEPs with nearly 70 aa to the *S. meliloti* annotation, that is, an increase in the number of annotated SEPs (present in at least one of the GenBank 2014 and RefSeq 2017 annotations) by approximately 17%.

Ribo-seq is a powerful technique for detecting translation on a global scale with high sensitivity (Ingolia et al. 2019). However, in contrast to eukaryotic model systems, codon resolution has not yet been achieved in Ribo-seq analyses of bacteria (Cianciulli Sesso et al. 2021; Mohammad et al. 2019; Vazquez-Laslop et al. 2022; Venturini et al. 2020). Trapping ribosomes on mRNA and generating ribosome footprints have remained challenging, requiring careful optimization for each bacterial species. Our Ribo-seq workflow for *S. meliloti* includes ribosome trapping by rapid cooling of the culture without using antibiotics and cell lysis in an adapted buffer, followed by digestion of unprotected RNA by RNase I, which is not inactivated by the ribosomes of *S. meliloti* (**Fig. 1B**). RNase I has the advantage of precisely cleaving at both 5′ and 3′ ends of ribosome-protected mRNA without sequence specificity, in contrast to the routinely used MNase (Bartholomäus et al. 2016). The digestion of 5′ and 3′ regions of translated mRNAs (**Fig. 1C**, **Fig. 3C**), higher TEs of annotated CDS in comparison to non-coding RNAs (**Fig. 2B**), and pronounced ribosome protection up to 16 nt upstream and downstream of start codons (**Fig. S2; Fig. 1C** and **1D**) show the successful establishment of Ribo-seq for *S. meliloti*.

In addition to providing the first genome-wide ribosome-binding map of a *Hyphomicrobiales* member, our Ribo-seq analysis uncovered translation for 85 annotated sORFs and identified 37 novel sORFs missing in the GenBank 2014 and Refseq 2017 annotations of the *S. meliloti* genome. The translated sORFs were found on all three replicons and had similar preferences for start and stop codons independently of whether they were annotated or novel (**Fig. 3** and **Fig. 4**). The novel sORFs were generally shorter than the annotated ones (**Fig. 4B; Table S4**), clearly showing the advantage of the Ribo-seq method for SEP discovery. Many of the novel sORFs were probably not annotated due to their location in short transcripts considered as non-coding RNAs or asRNAs or in 5′ and 3′ UTRs (**Fig. 4C**).

Several translated novel sORFs internal to annotated genes (nested ORFs; Gray et al. 2022) were also predicted by our Ribo-seq data. However, they were excluded from the analysis as additional evidence is needed to confirm their existence. Targeted detection of translation initiation sites is useful in uncovering such sORFs by Ribo-seq (Meydan et al. 2019; Weaver et al. 2019), a strategy beyond the scope of our study. However, the existence of an internal sORF predicted by Ribo-seq was supported by the MS detection of a novel, 34-aa-long SEP translated in a different frame in the genomic region encoding SM2011_b20335 (**Fig. 5C** and **Fig. S5G;** sORF59 in **Fig. 7**).

A challenge in defining novel sORFs for any genome is that annotations from different reference genome annotation centers can differ substantially for an identical sequence and change over time; that is, CDS are being added but are also removed in more recent annotations (see **Fig. 2F** and the “master” **Table S4**). Accordingly, two of the 48 novel SEPs are now bona fide-predicted CDS in the latest RefSeq 2022 annotation, with MS-evidence of a single PSM found with the custom iPtgxDB, whereas two other Ribo-seq-identified sORFs have variable pseudogene status in different annotation releases (see **Table S4**). iPtgxDBs, which integrate existing reference annotations and add *in silico* predicted ORFs in all six frames to virtually cover the entire protein coding potential of a prokaryote, can be used to overcome such problems and enable MS-based detection of novel SEPs (Omasits et al. 2017).

Here, in addition to a standard large iPtgxDB of *S. meliloti* (Melior et al. 2020), we applied the concept of a small, custom iPtgxDB lacking *in silico* predictions and including the top predictions from our experimental Ribo-seq data. This custom iPtgxDB is approximately 20-fold smaller and benefits statistics and FDR estimation (Li et al. 2016; Blakeley et al. 2012). Notably, although the identification of 11 *in silico* predicted novel sORFs was possible only with the standard iPtgxDB, the small iPtxDB contributed substantially to the validation of annotated sORFs, increasing the number of SEPs with experimental support by 10% (**Fig. S4**). The detection of more SEPs was also facilitated by applying three experimental approaches, two of which included enrichment of small proteins. The MS detection of enriched SEPs without a proteolytic digest, including, for example, the 12 aa proteolysis tag encoded by tmRNA (**Fig. 5C** and **Fig. S5**), shows that this method can be useful for the identification of SEPs.

The validation of translation by Western blot analysis for 15 out of 20 analyzed novel SEPs with Ribo-seq support (**Fig. 6B–6F; Table S7**), only three of which were detected by MS with at least 2 PSMs (**Table S4**), underlines the power of the Ribo-seq technique for identification of translated sORFs. The example of SEP7 (**Fig. 6B**; 59 aa, restriction endonuclease-like, conserved in *Rhizobiaceae*), which was added to the RefSeq annotation 2017 and was detected by 2 PSMs using the small, custom iPtgxDB, illustrates the added value of the latter. Detection of translation for the novel SEP1 (23 aa, conserved in *Rhizobiaceae*) by Western blot analysis (**Fig. 6E**) and Ribo-seq (highest TE among the non-annotated translated sORFs, **Table S7**), even though it was identified by only 1 PSM in the MS analysis (**Fig. S5E**), suggests that putative SEPs with 1 PSM can be truly expressed, real small proteins. Similarly, the annotated SEP20 (46 aa, conserved in Alphaproteobacteria) was confirmed by Western blot analysis (**Fig. 6G**), although it had only 1 PSM (**Fig. S5F**) and did not pass the stringent manual evaluation of the Ribo-seq data (**Table S7**). We suggest that the conservation analysis of putative SEPs, which have minimal MS evidence (e.g., 1 PSM) and/or correspond to sORFs that did not pass the very stringent manual curation of the Ribo-seq data, can help define SEP candidates with potentially important functions that can be validated and analyzed in the future.

Despite the lower sensitivity of MS compared with Ribo-seq, using MS we detected 16 additional SEPs that were not identified as translated by Ribo-seq. Eleven of them were novel, showing the importance of complementary methods for comprehensive analysis of bacterial small proteomes. The reported numbers of validated and novel sORFs and their encoded SEPs are affected by the somewhat arbitrary cut-off of 70 aa. In fact, our data provide evidence for the translation of three additional, non-annotated proteins below 100 aa, which are considered small in other studies (Baumgartner et al. 2016; Kaulich et al. 2021; VanOrsdel et al. 2018) (see **Table S4**).

The functions of small proteins are difficult to predict *in silico*, often because they are too small to harbor known protein domains or motifs (Ahrens et al. 2022). In addition, for SEPs smaller than 30 aa *in silico* analysis by Phyre2 is still impossible. Keeping these limitations in mind, we present a list of putative functions corresponding to Phyre2 best hits (**Table S8**). Since modeling of a partial SEP sequence by Phyre2 may provide a hint of potential interactions with other proteins or protein complexes, we mention predictions based on greater than 30% confidence homology in **Fig. 7**, including the predicted DNA-binding function of the 43 aa SEP38 and a potential role in bleomycin resistance of the 39 aa SEP34. SEP function can also be predicted based on gene synteny (Ahrens et al. 2022), as mentioned above, for SEP5 (potential leader peptide encoded by a regulatory upstream sORF) and SEP20 encoded in a cytochrome oxidase operon. Our findings show that, excluding the tmRNA sORF, 13 out of the 48 novel SEPs (sORFs) are conserved in *Rhizobiaceae*, seven in *Hyphomicrobiales*, and three in at least two bacterial phyla, which likely suggests physiological relevance. Most of the translated sORFs or SEPs were detected in logarithmic cultures grown in a minimal medium, where bacteria synthesize virtually all metabolites for cell reproduction. Thus, some of these SEPs can be of general importance for growth or are needed for survival and competitiveness under oligotrophic conditions in soil and rhizosphere.

In summary, our work shows that a combination of methods is beneficial for increasing the number of experimentally validated SEPs. Using Ribo-seq, MS, and Western blot analysis of C-terminally tagged proteins, we provide evidence for the translation of 48 SEPs with ≤ 70 aa to be added to the annotation of *S. meliloti*, thus substantially increasing the number of cataloged SEPs. With the MS data, the corresponding full and small custom iPtgxDBs, and importantly, the first Ribo-seq analysis of a *Hyphomicrobiales* member, which can be viewed with an interactive online JBrowse instance (http://www.bioinf.uni-freiburg.de/ribobase), our study provides valuable resources for future studies on and beyond the small proteome.

## ACKNOWLEDGEMENTS

We thank Stephanie Färber for the technical assistance in growing *S. meliloti* to establish Ribo-seq, and Pierre-Alexander Mücke, as well as Claudia Hirschfeld, for MS measurements. The cell-free lysates for the MS analysis were provided by Hendrik Melior. We thank Sarah L. Svensson for the critical feedback on the manuscript. This work was supported by the German Research Foundation (DFG) priority program SPP2002 “Small Proteins in Prokaryotes, an Unexplored World” grants (grant SH580/7-1 and SH580/7-2 to CMS, grant BA 2168/21-2 to RB, grant BE 3869/5-1 and BE 3869/5-2 to DB, and grant Ev 42/7-1 to EEH). Additional funding was received from the Swiss National Science Foundation (grant 197391) to CHA and from DFG (GRK2355 project number 325443116) to EEH and Germany’s Excellence Strategy (CIBSS – EXC-2189 – Project ID 390939984) to RB. Computational resources were provided by the BMBF-funded de.NBI Cloud within the German Network for Bioinformatics Infrastructure (de.NBI) (031A532B, 031A533A, 031A533B, 031A534A, 031A535A, 031A537A, 031A537B, 031A537C, 031A537D, 031A538A).

## AUTHOR CONTRIBUTIONS

CMS, RB, and EEH initiated the project. LH, RG, SM, RS, SA, CHA, BH, and EEH designed the experiments and analyzed the data. LH established Ribo-seq for *S. meliloti,* performed the Ribo-seq analysis, manually examined the implied novel sORFs, explored evolutionary conservation, and predicted the function of novel SEPs. RG performed the bioinformatic processing of the Ribo-seq data. SM conducted the mass-spectrometry experiments, database searches and manually evaluated spectra implying novel SEPs. BH and CHA contributed to the proteogenomic analysis to identify the known and novel SEPs, explored the value of iPtgxDBs for Ribo-seq data, consolidated experimental results in a master table, and identified annotation differences. RS, SA, and SBW performed the cloning and Western blot analyses. LH, CHA, and EEH mainly wrote the manuscript with input and feedback from the other authors. DB, RB, CMS, CHA, and EEH supervised the research and provided resources and funding. All authors approved the submitted version.

## List of Supplementary Figures and Supplementary Tables

**Figure S1. Schematic representation of the plasmids used in this study**. pRS1: The plasmid that was used as a vector backbone (the multiple cloning site [MCS] is shown) to create the following plasmids (all shown below): pRS1-T*rrn*: RS1 derivative with cloned transcriptional terminator (T*rrn*). pSW1: pRS1-T*rrn* derivative with cloned sequence of the SPA-tag (SPA), preceded by a linker (L). pSW2: pSW1 derivative with cloned promoter P*sinI*, which is constitutively active in *Sinorhizobium meliloti* 2011 during growth. pSW2-SEP: pSW2 derivative with a cloned sORF and its −15 5′ UTR region potentially harboring a Shine-Dalgarno sequence; the sORF is cloned without the stop codon in frame with the linker and the SPA tag.

**Figure S2. Metagene analysis of *Sinorhizobium meliloti* ribosome footprints.** Genome-wide analysis of ribosome occupancy near annotated start codons. The recovered ribosome footprints (length distribution varies from 27 nt to 33 nt) were mapped using **(A)** 5′ end and **(B)** 3′ end approaches. Metagene analysis of the 32-nt-long ribosome footprints by the 5′ end **(C)** and 3′ end **(D)** mapping approaches shows that the ribosome protects a region of 16 nt upstream and downstream of annotated start codons (+1; first nucleotide of the start codon).

**Figure S3. Examples of genomic regions with high translation efficiency (TE). (A)** The *gabD1* gene (SM2011_c02780) encoding for succinate semialdehyde dehydrogenase harbors a short 5′ UTR (24 nt), which exhibits ribosome protection at −15/−16 upstream of the start codon that contributes to its high TE value (TE = 34.6). **(B)** The 3′ UTR of the SM2011_c01202 gene encoding lipoprotein shows high ribosome density and TE. HRIBO predicts a potential novel downstream small open reading frame (sORF) in this region (25 amino acids [aa], TE = 4.3). **(C)** A novel sORF is predicted in the non-coding small RNA (sRNA) SMc06505 (38 aa, TE = 3.23). This can be an example of dual-function sRNA in *Sinorhizobium meliloti* 2011. Genomic locations and coding regions are indicated below the image. Bent arrows indicate the transcription start sites based on (Sallet et al. 2013).

**Figure S4. Comparison of search results against the standard and custom-integrated proteogenomic search databases (iPtgxDBs). (A)** Venn diagram with an overview of the proteins identified by the three different experimental approaches (the colors match those from Figure 5A: gray, green, and blue represent the trypsin digest, SPE and Lys-C digest, and SPE and no protease digestion, respectively). **(B)** The Venn diagram shows the overlap of the number of proteins identified in the searches against the two iPtgxDBs (standard and custom iPtgxDB). The search against the 20-fold smaller custom iPtgxDb allowed the identification of 112 proteins that were not identified in the search against the much larger standard iPtgxDB. These hits include RefSeq or GenBank annotations. The 18 unique identifications made with the standard iPtgxDB include novel proteins or proteoforms contributed from Chemgenome *ab initio* predictions or *in silico* predictions that are not contained in the small custom iPtgxDB.

**Figure S5. Mass spectrometry of selected novel small open reading frame-encoded proteins (SEPs).** Here, we show some of the spectra that allowed us to identify novel SEPs. If more than one peptide spectrum match (PSM) was detected for a given peptide ion, a representative spectrum was selected. MS2 spectra with assigned fragment ion m/z (left) and fragmentation tables (right) were obtained with Scaffold V4.8.7 using the search output files (*.sf3), which were deposited at the ProteomeXchange Consortium with the dataset identifier PXD034931. Colored m/z were assigned in the identifying MS2 spectra.

**Figure S6. Analysis of a putative small open reading frame (sORF) in tmRNA of *Sinorhizobium meliloti* 2011. (A)** Schematic view of the chromosomal *ssrA* locus corresponding to tmRNA, which is discontinuous in Alphaproteobacteria due to post-transcriptional removal of the indicated internal segment (Keiler et al. 2000; Ulvé et al. 2007). The tRNA and mRNA parts of the tmRNA and their lengths are indicated. In the mRNA part, a putative sORF corresponding to a 23-amino acid (aa) SEP was predicted (the potential SEP sequence is shown). The 3′ part of the sORF corresponds to the proteolytic tag-encoding sequence (the alanine encoded by the resume codon is shown in bold). The indicated parts of the *ssrA* gene were cloned in pSW2, and the *S. meliloti* 2011 strains containing the corresponding plasmids were used for Western blot analysis. While no specific bands were detected in lysates of the pSW2-ssrA1-containing plasmid, which lacks the tRNA part of tmRNA (data not shown), a strong signal was obtained with the pSW2-ssrA2 plasmid, which contains both the tRNA and mRNA parts (see panel C). **(B)** Schematic representation of the *ssrA* part cloned in pSW2-ssrA2. The used mutations are indicated. 1: Conserved GG nucleotides upstream of the putative start codon TTG were mutated to TT. 2: The resume codon was changed to encode valine instead of alanine. 3: The putative start codon TTG was mutated to the stop codon TAG. **(C)** Western blot analysis with antibodies directed against the FLAG part of the sequential peptide affinity (SPA) tag, which was fused in frame to the proteolytic tag. Crude lysates (corresponding to 20 OD) of *S. meliloti* 2011 strains containing the indicated plasmids (see panels A and B) were analyzed. The detected bands above 25 kDA have identical lengths. Expression of the corresponding SPA-tagged peptide was abolished by the indicated GG/TT mutation, which disrupts conserved base pairing in the tmRNA (Keiler et al. 2000), whereas a weak signal was still detected when the resume codon was mutated. Destroying the putative 23 aa sORF (TTG/TAG mutation in variant 3) did not abolish the expression of the tagged peptide. These results, combined with no detection of a peptide using pSW2-ssrA1, suggest that the SPA-tagged peptide detected in panel C corresponds to the 12 aa proteolytic tag and not to the putative 23 aa SEP. Furthermore, the data support the important role of the analyzed GG nucleotides for the tmRNA function.

**Figure S7**. **Analysis of the S100 and P100 fractions of *Sinorhizobium meliloti* 2011 strains producing the indicated small open reading frame-encoded proteins (SEPs) from pSW2-SEP plasmids.** The empty vector control (EVC) strain was used as the negative control. Identical volumes of the S100 and P100 fractions were loaded. Top panels: Western blot analysis with monoclonal anti-FLAG antibodies. Bottom panels: Coomassie-stained gels, in which the used protein fractions are shown. Migration of marker proteins (in kDa) is shown on the left side, and exposition times are indicated. * Unspecific signal.

## Supplementary Tables

**Table S1. Distribution of total mapped reads to different annotated RNA classes.**

The table indicates the total number of mapped reads mapping to annotations for different RNA classes (mRNA, rRNA, and tRNA).

**Table S2. Translation efficiency values for all annotated genes of *Sinorhizobium meliloti* 2011.**

This table is generated by the HRIBO pipeline and contains information about the annotated features extracted from the used GenBank 2014 annotation. It contains information, such as the gene identifier, the start and stop codons, the encoding strand, locus tags, nucleotides/amino acid sequences, and information regarding the encoded protein, translation efficiency (TE), as well as the RPKM values. The RPKM values were calculated directly after the removal of multi-mapping reads and before the removal of reads mapping to rRNA.

**Table S3. Overview of the six integrated proteogenomic search databases (iPtgxDBs) created for this project.**

Three large standard iPtgxDBs were created: one for trypsin, one for Lys-C, and a modified version for the experimental approach without proteolytic digestion. On the top, three approximately 20-fold smaller custom iPtgxDBs were created using experimental ribosome profiling data instead of the Chemgenome *ab initio* predictions and the large number of *in silico* predictions that aim to capture the entire protein coding potential.

**Table S3.1:** Size of the six annotation sources (three reference genome annotations, two *ab initio* predictions, and the *in silico* prediction) and the added novelty (new clusters, new reductions, and new extensions) achieved through the step-wise, hierarchical integration carried out for the standard iPtgxDBs.

**Table S3.2:** Database entries and number of excluded entries for the three standard iPtgxDBs for trypsin, Lys-C, and without proteolytic digestion.

**Table S3.3:** Size of the five annotation sources (three reference genome annotations, one *ab initio* prediction, and experimental ribosome profiling data) and the added novelty (new clusters, new reductions, new extensions) achieved through the step-wise, hierarchical integration carried out for the three smaller custom iPtgxDBs.

**Table S3.4:** Database entries and number of excluded entries for the three custom iPtgxDBs for trypsin, LysC, and without proteolytic digestion.

**Table S4. “Master table”, with an overview of various datasets detailed in the manuscript.**

This table lists several datasets described in the manuscript and allows researchers to select/filter them. It contains information about genomic location and annotation for the 6263 CDS (GenBank 2014 annotation) and their match with the RefSeq 2017 annotation. For the ribosome profiling (Ribo-seq) data, we listed the 3,758 GenBank annotated open reading frames (ORFs) identified as translated in minimal medium (translation efficiency [TE] of ≥ 0.5 and RNA-seq and Ribo-seq RPKM of ≥ 10), all 259 GenBank annotated small (sORFs) (≤ 70 amino acids [aa]), and the 85 GenBank annotated sORFs identified as translated after manual curation (Figures 2 and 3). Furthermore, listed are the 266 potentially novel Ribo-seq implied sORFs (not present in the GenBank 2014 annotation; TE of ≥ 0.5, RNA-seq and Ribo-seq RPKM of ≥ 10, DeepRibo score of > −0.5), and the subset of 54 manually curated, high-confidence non-annotated sORFs (Figure 4). For the mass spectrometry (MS) data, a list of 191 sORF-encoded proteins (SEPs) can be filtered, among which at least one peptide spectrum match (PSM) was identified, the 58 SEPs that pass the more stringent PSM filters (Figure 5), and the respective search database results where they were detected (standard integrated proteogenomic search databases [iPtgxDB], custom iPtgxDB or both). A total of 20 candidates for which Western blot validation was attempted (15 thereof with success) can then be selected (Figure 6), as well as the 48 novel sORFs missing from both the GenBank 2014 and Refseq 2017 annotations, and 37 detected by Ribo-seq and 11 by proteomics (Figure 7). Finally, the PSM counts and identifiers contained in the large standard iPtgxDB and the small custom iPtgxDB of the 1,219 proteins identified by MS are provided.

**Table S5. Oligonucleotides used in this study.**

**Table S6**. **List of annotated open reading frames (ORFs) and small ORFs (sORFs) detected as translated in *Sinorhizobium meliloti* 2011 by ribosome profiling (Ribo-seq).**

Using Ribo-seq (average translation efficiency [TE] of ≥ 0.5 and RNA-seq and Ribo-seq RPKM of ≥ 10 in both replicates), translation was detected for 3,758 ORFs out of the 6,263 annotated ORFs. In addition, we manually curated each annotated sORF with an average TE of ≥ 0.5 and RNA-seq and Ribo-seq RPKM of ≥ 10 (in both replicates) in a genome browser. Based on the manual curation, translation was detected for 85 of the 259 annotated *Sinorhizobium meliloti* sORFs (GenBank annotation 2014).

**Table S7: List of non-annotated translated small open reading frames (sORFs) identified in *Sinorhizobium meliloti* 2011 by ribosome profiling (Ribo-seq).**

The two prediction tools DeepRibo and Reparation, which are implemented in the HRIBO pipeline, predicted a high number of translated sORFs (approximately 15,000 sORFs). To cope with and prioritize this high number of predictions, we first applied cut-offs (average translation efficiency of ≥ 0.5 and RNA-seq and Ribo-seq RPKM of ≥ 10 in both replicates, and DeepRibo score of > −0.5), which resulted in a list of 266 potential, translated, non-annotated sORFs (see Table S4) (GenBank annotation 2014). We then manually curated each of these 266 predicted sORFs in a genome browser. The resulting list of 54 non-annotated sORFs, which we propose with high confidence to be translated, is presented in this table..

**Table S8: List of the 48 novel small open reading frames identified in *Sinorhizobium meliloti* 2011 using ribosome profiling and proteomics.** A list of Phyre2 best hits is also provided.

