## Supplemental Figures 1 to 7 for "Small proteome of the nitrogen-fixing plant symbiont *Sinorhizobium meliloti*"

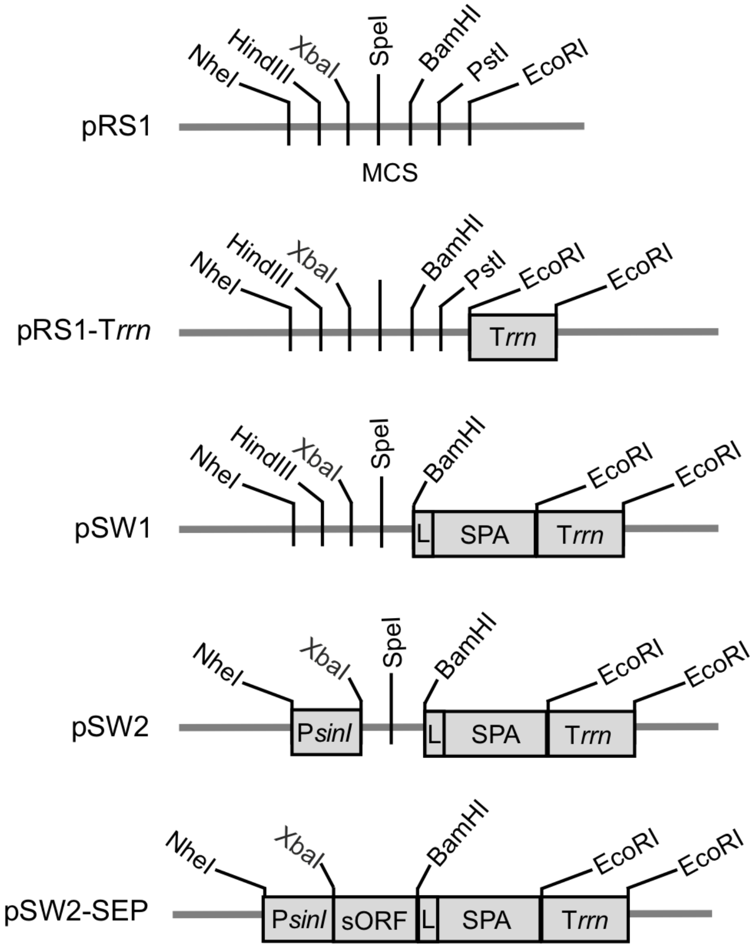


**Figure S1. Schematic representation of the plasmids used in this study**. pRS1: The plasmid that was used as a vector backbone (the multiple cloning site [MCS] is shown) to create the following plasmids (all shown below): pRS1-$T_{rrn}$: RS1 derivative with cloned transcriptional terminator ($T_{rrn}$). pSW1: pRS1-$T_{rrn}$ derivative with cloned sequence of the SPA-tag (SPA), preceded by a linker (L). pSW2: pSW1 derivative with cloned promoter $P_{sinI}$, which is constitutively active in *Sinorhizobium meliloti* 2011 during growth. pSW2-SEP: pSW2 derivative with a cloned sORF and its −15 5′ UTR region potentially harboring a Shine-Dalgarno sequence; the sORF is cloned without the stop codon in frame with the linker and the SPA tag.

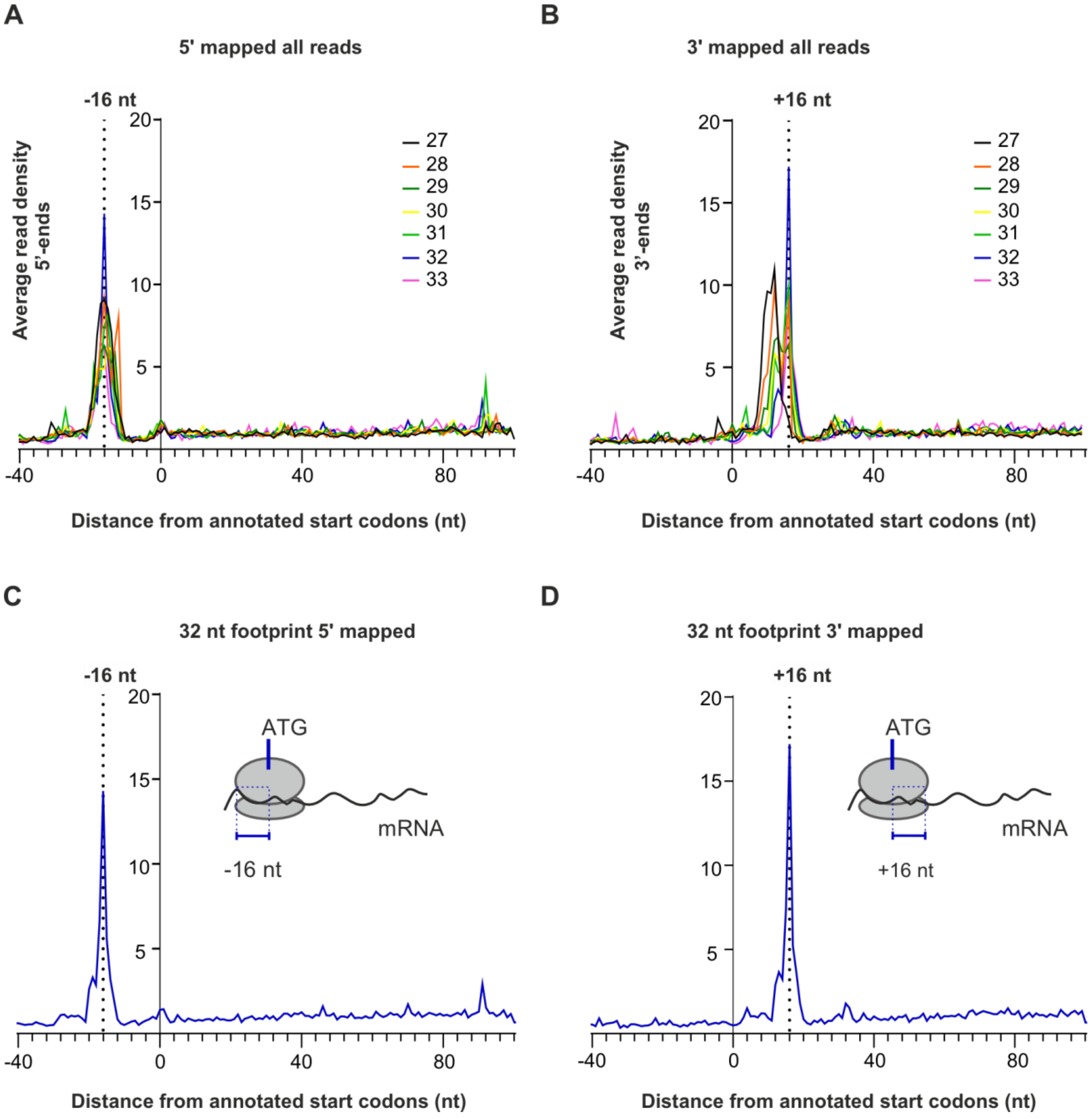


**Figure S2. Metagene analysis of *Sinorhizobium meliloti* ribosome footprints.** Genome-wide analysis of ribosome occupancy near annotated start codons. The recovered ribosome footprints (length distribution varies from 27 nt to 33 nt) were mapped using **(A)** 5′ end and **(B)** 3′ end approaches. Metagene analysis of the 32-nt-long ribosome footprints by the 5′ end **(C)** and 3′ end **(D)** mapping approaches shows that the ribosome protects a region of 16 nt upstream and downstream of annotated start codons (+1; first nucleotide of the start codon).

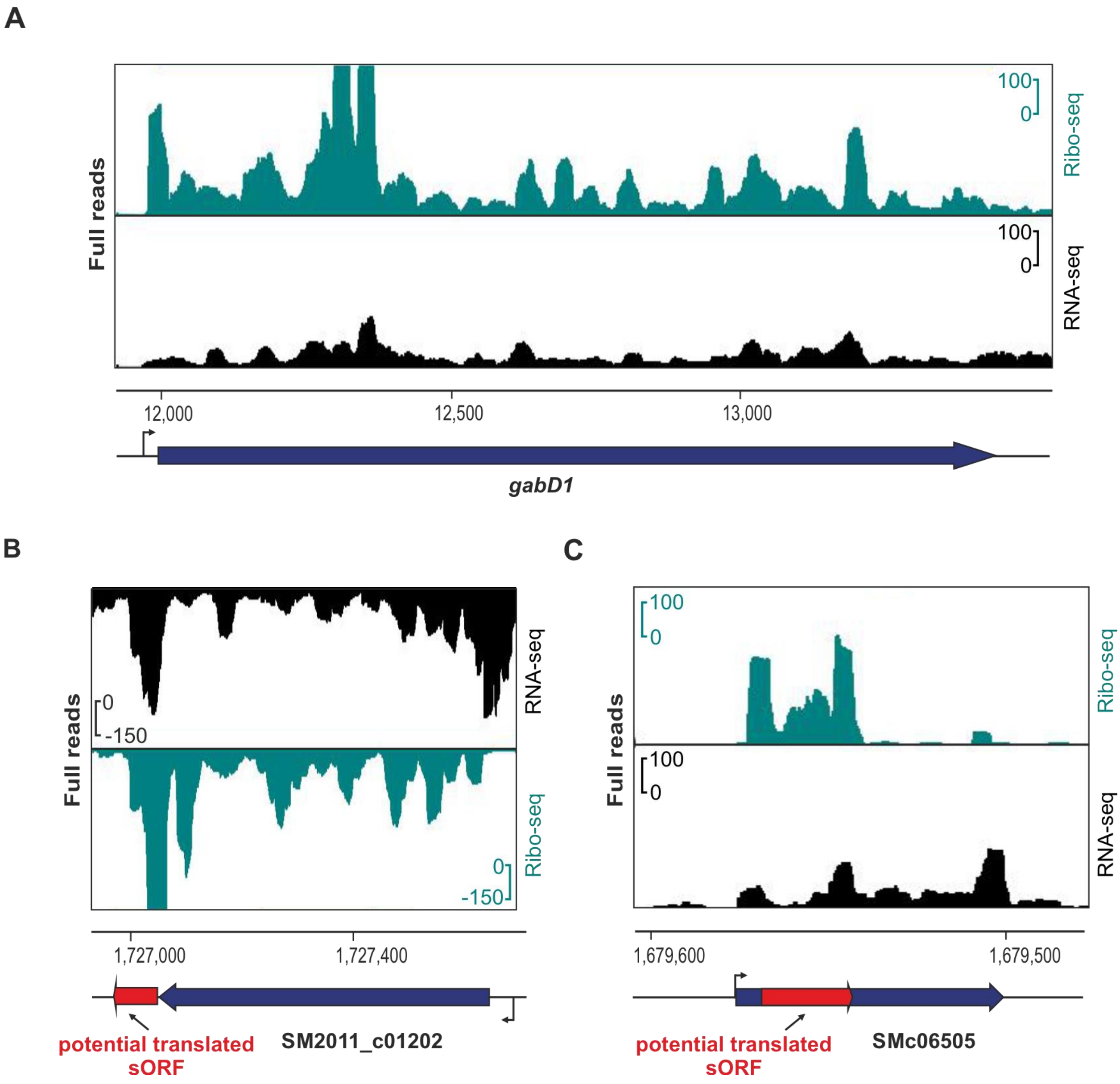


**Figure S3. Examples of genomic regions with high translation efficiency (TE). (A)** The *gabD1* gene (SM2011_c02780) encoding for succinate semialdehyde dehydrogenase harbors a short 5′ UTR (24 nt), which exhibits ribosome protection at −15/−16 upstream of the start codon that contributes to its high TE value (TE = 34.6). **(B)** The 3′ UTR of the SM2011_c01202 gene encoding lipoprotein shows high ribosome density and TE. HRIBO predicts a potential novel downstream small open reading frame (sORF) in this region (25 amino acids [aa], TE = 4.3). **(C)** A novel sORF is predicted in the non-coding small RNA (sRNA) SMc06505 (38 aa, TE = 3.23). This can be an example of dual-function sRNA in *Sinorhizobium meliloti* 2011. Genomic locations and coding regions are indicated below the image. Bent arrows indicate the transcription start sites based on (Sallet et al. 2013)**.**

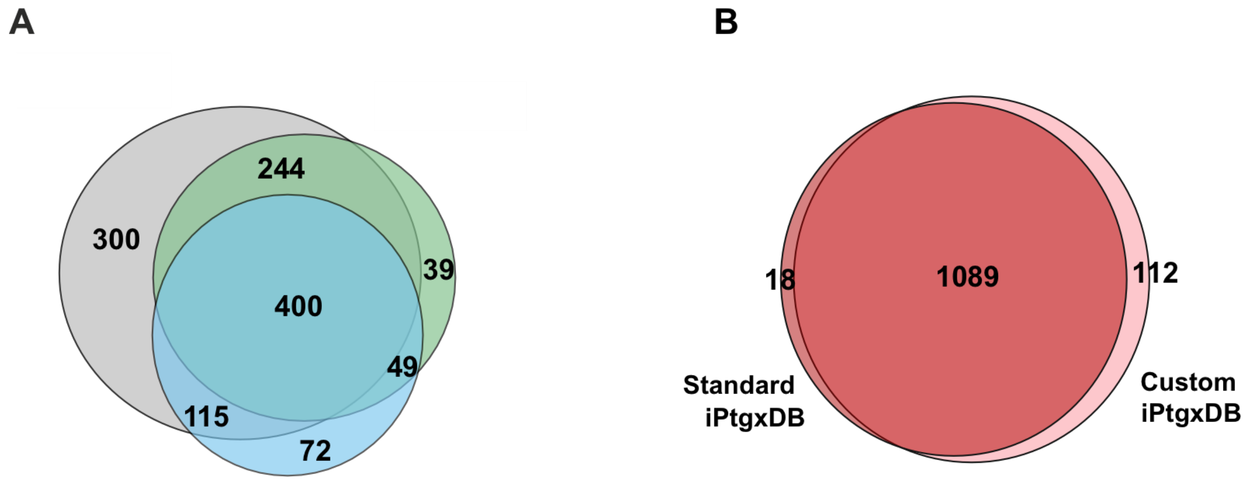


**Figure S4. Comparison of search results against the standard and custom-integrated proteogenomic search databases (iPtgxDBs). (A)** Venn diagram with an overview of the proteins identified by the three different experimental approaches (the colors match those from Figure 5A: gray, green, and blue represent the trypsin digest, SPE and Lys-C digest, and SPE and no protease digestion, respectively). **(B)** The Venn diagram shows the overlap of the number of proteins identified in the searches against the two iPtgxDBs (standard and custom iPtgxDB). The search against the 20-fold smaller custom iPtgxDb allowed the identification of 112 proteins that were not identified in the search against the much larger standard iPtgxDB. These hits include RefSeq or GenBank annotations. The 18 unique identifications made with the standard iPtgxDB include novel proteins or proteoforms contributed from Chemgenome *ab initio* predictions or *in silico* predictions that are not contained in the small custom iPtgxDB.

**Fig. S5A**
**Spectra to Fig. 5C. SEP identifier in Table S4: CP004140.1:2615856-2616038:+; 14 aa extension of HmuP**

1 Peptide of CP004140.1:2615856-2616038:+, 2 PSM with standard iPtgxDB in MM after tryptic digest of non-enriched proteins

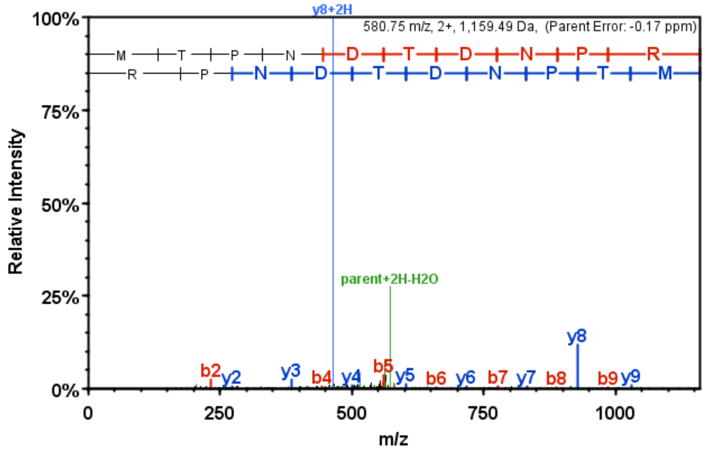


| B | B Ions | B+2H | B-NH3 | B-H2O | AA | Y Ions | Y+2H | Y-NH3 | Y-H2O | Y |
|---|---|---|---|---|---|---|---|---|---|---|
| 1 | 132.0 | | | | M | 1,160.5 | 580.8 | 1,143.5 | 1,142.5 | 10 |
| 2 | 233.1 | | | 215.1 | T | 1,029.5 | 515.2 | 1,012.4 | 1,011.4 | 9 |
| 3 | 330.1 | | | 312.1 | P | 928.4 | 464.7 | 911.4 | 910.4 | 8 |
| 4 | 444.2 | | 427.2 | 426.2 | N | 831.4 | 416.2 | 814.3 | 813.3 | 7 |
| 5 | 559.2 | | 542.2 | 541.2 | D | 717.3 | 359.2 | 700.3 | 699.3 | 6 |
| 6 | 660.3 | 330.6 | 643.2 | 642.3 | T | 602.3 | | 585.3 | 584.3 | 5 |
| 7 | 775.3 | 388.2 | 758.3 | 757.3 | D | 501.2 | | 484.2 | 483.2 | 4 |
| 8 | 889.3 | 445.2 | 872.3 | 871.3 | N | 386.2 | | 369.2 | | 3 |
| 9 | 986.4 | 493.7 | 969.4 | 968.4 | P | 272.2 | | 255.1 | | 2 |
| 10 | 1,160.5 | 580.8 | 1,143.5 | 1,142.5 | R | 175.1 | | 158.1 | | 1 |

2 Peptides of CP004140.1:2615856-2616038:+, 1 PSM each with standard iPtgxDB in MM after SPE-enrichment, without proteolytic digest

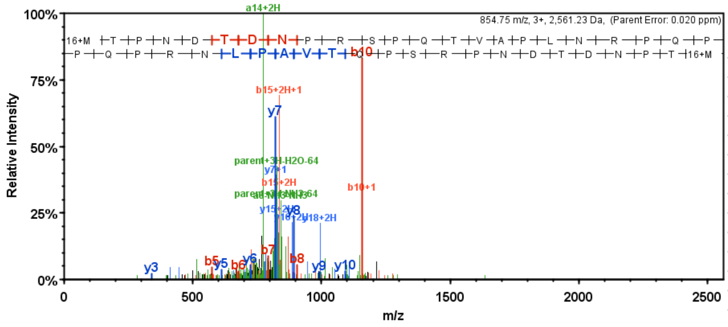


| B | B Ions | B+2H | B-NH3 | B-H2O | AA | Y Ions | Y+2H | Y-NH3 | Y-H2O | Y |
|---|---|---|---|---|---|---|---|---|---|---|
| 1 | 148.0 | 74.5 | | | M+16 | 2,562.2 | 1,281.6 | 2,545.2 | 2,544.2 | 23 |
| 2 | 249.1 | 125.0 | | 231.1 | T | 2,415.2 | 1,208.1 | 2,398.2 | 2,397.2 | 22 |
| 3 | 346.1 | 173.6 | | 328.1 | P | 2,314.2 | 1,157.6 | 2,297.1 | 2,296.1 | 21 |
| 4 | 460.2 | 230.6 | 443.2 | 442.2 | N | 2,217.1 | 1,109.1 | 2,200.1 | 2,199.1 | 20 |
| 5 | 575.2 | 288.1 | 558.2 | 557.2 | D | 2,103.1 | 1,052.0 | 2,086.0 | 2,085.0 | 19 |
| 6 | 676.3 | 338.6 | 659.2 | 658.3 | T | 1,988.0 | 994.5 | 1,971.0 | 1,970.0 | 18 |
| 7 | 791.3 | 396.1 | 774.3 | 773.3 | D | 1,887.0 | 944.0 | 1,870.0 | 1,869.0 | 17 |
| 8 | 905.3 | 453.2 | 888.3 | 887.3 | N | 1,772.0 | 886.5 | 1,754.9 | 1,753.9 | 16 |
| 9 | 1,002.4 | 501.7 | 985.4 | 984.4 | P | 1,657.9 | 829.5 | 1,640.9 | 1,639.9 | 15 |
| 10 | 1,158.5 | 579.7 | 1,141.5 | 1,140.5 | R | 1,560.9 | 780.9 | 1,543.8 | 1,542.9 | 14 |
| 11 | 1,245.5 | 623.3 | 1,228.5 | 1,227.5 | S | 1,404.8 | 702.9 | 1,387.7 | 1,386.7 | 13 |
| 12 | 1,342.6 | 671.8 | 1,325.5 | 1,324.6 | P | 1,317.7 | 659.4 | 1,300.7 | 1,299.7 | 12 |
| 13 | 1,470.6 | 735.8 | 1,453.6 | 1,452.6 | Q | 1,220.7 | 610.8 | 1,203.6 | 1,202.7 | 11 |
| 14 | 1,571.7 | 786.3 | 1,554.6 | 1,553.7 | T | 1,092.6 | 546.8 | 1,075.6 | 1,074.6 | 10 |
| 15 | 1,670.7 | 835.9 | 1,653.7 | 1,652.7 | V | 991.6 | 496.3 | 974.5 | | 9 |
| 16 | 1,741.8 | 871.4 | 1,724.8 | 1,723.8 | A | 892.5 | 446.8 | 875.5 | | 8 |
| 17 | 1,838.8 | 919.9 | 1,821.8 | 1,820.8 | P | 821.5 | 411.2 | 804.4 | | 7 |
| 18 | 1,951.9 | 976.5 | 1,934.9 | 1,933.9 | L | 724.4 | 362.7 | 707.4 | | 6 |
| 19 | 2,066.0 | 1,033.5 | 2,048.9 | 2,048.0 | N | 611.3 | 306.2 | 594.3 | | 5 |
| 20 | 2,222.1 | 1,111.5 | 2,205.0 | 2,204.1 | R | 497.3 | 249.1 | 480.3 | | 4 |
| 21 | 2,319.1 | 1,160.1 | 2,302.1 | 2,301.1 | P | 341.2 | 171.1 | 324.2 | | 3 |
| 22 | 2,447.2 | 1,224.1 | 2,430.1 | 2,429.2 | Q | 244.1 | 122.6 | 227.1 | | 2 |
| 23 | 2,562.2 | 1,281.6 | 2,545.2 | 2,544.2 | P | 116.1 | 58.5 | | | 1 |

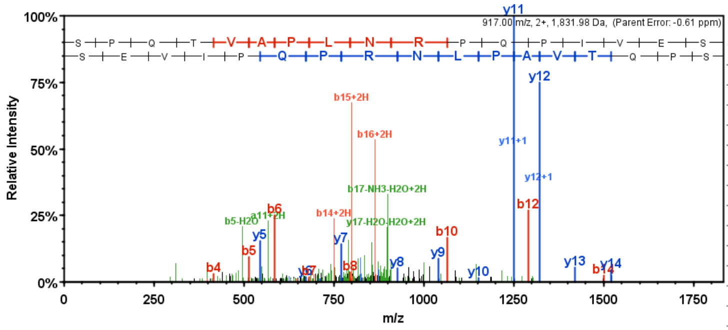


| B | B Ions | B+2H | B-NH3 | B-H2O | AA | Y Ions | Y+2H | Y-NH3 | Y-H2O | Y |
|---|---|---|---|---|---|---|---|---|---|---|
| 1 | 88.0 | | | 70.0 | S | 1,833.0 | 917.0 | 1,816.0 | 1,815.0 | 17 |
| 2 | 185.1 | | | 167.1 | P | 1,746.0 | 873.5 | 1,728.9 | 1,727.9 | 16 |
| 3 | 313.2 | | 296.1 | 295.1 | Q | 1,648.9 | 825.0 | 1,631.9 | 1,630.9 | 15 |
| 4 | 414.2 | | 397.2 | 396.2 | T | 1,520.8 | 760.9 | 1,503.8 | 1,502.8 | 14 |
| 5 | 513.3 | | 496.2 | 495.3 | V | 1,419.8 | 710.4 | 1,402.8 | 1,401.8 | 13 |
| 6 | 584.3 | 292.7 | 567.3 | 566.3 | A | 1,320.7 | 660.9 | 1,303.7 | 1,302.7 | 12 |
| 7 | 681.4 | 341.2 | 664.3 | 663.3 | P | 1,249.7 | 625.3 | 1,232.7 | 1,231.7 | 11 |
| 8 | 794.4 | 397.7 | 777.4 | 776.4 | L | 1,152.6 | 576.8 | 1,135.6 | 1,134.6 | 10 |
| 9 | 908.5 | 454.7 | 891.5 | 890.5 | N | 1,039.6 | 520.3 | 1,022.5 | 1,021.5 | 9 |
| 10 | 1,064.6 | 532.8 | 1,047.6 | 1,046.6 | R | 925.5 | 463.3 | 908.5 | 907.5 | 8 |
| 11 | 1,161.6 | 581.3 | 1,144.6 | 1,143.6 | P | 769.4 | | 752.4 | 751.4 | 7 |
| 12 | 1,289.7 | 645.4 | 1,272.7 | 1,271.7 | Q | 672.4 | | 655.3 | 654.3 | 6 |
| 13 | 1,386.7 | 693.9 | 1,369.7 | 1,368.7 | P | 544.3 | | | 526.3 | 5 |
| 14 | 1,499.8 | 750.4 | 1,482.8 | 1,481.8 | I | 447.2 | | | 429.2 | 4 |
| 15 | 1,598.9 | 800.0 | 1,581.9 | 1,580.9 | V | 334.2 | | | 316.2 | 3 |
| 16 | 1,727.9 | 864.5 | 1,710.9 | 1,709.9 | E | 235.1 | | | 217.1 | 2 |
| 17 | 1,833.0 | 917.0 | 1,816.0 | 1,815.0 | S | 106.0 | | | 88.0 | 1 |

See next page

**Fig. S5B**

**Spectra to Fig. 5C. SEP identifier in Table S4: CP004140.1:2290880-2290951:+; tmRNA-encoded proteolytic tag**

4 Peptides of CP004140.1:2290880-2290951:+ with 8 PSMs with standard iPtgxDB in MM after SPE-enrichment, without proteolytic digest

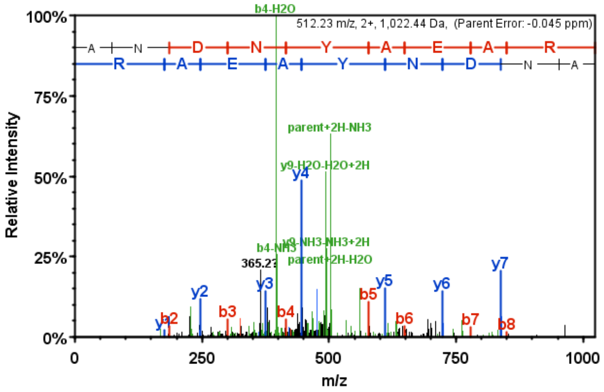


| B | B Ions | B+2H | B-NH3 | B-H2O | AA | Y Ions | Y+2H | Y-NH3 | Y-H2O | Y |
|---|---|---|---|---|---|---|---|---|---|---|
| 1 | 72.0 | | | | A | 1,023.4 | 512.2 | 1,006.4 | 1,005.4 | 9 |
| 2 | 186.1 | | 169.1 | | N | 952.4 | 476.7 | 935.4 | 934.4 | 8 |
| 3 | 301.1 | | 284.1 | 283.1 | D | 838.4 | 419.7 | 821.3 | 820.4 | 7 |
| 4 | 415.2 | | 398.1 | 397.1 | N | 723.3 | 362.2 | 706.3 | 705.3 | 6 |
| 5 | 578.2 | | 561.2 | 560.2 | Y | 609.3 | | 592.3 | 591.3 | 5 |
| 6 | 649.3 | 325.1 | 632.2 | 631.2 | A | 446.2 | | 429.2 | 428.2 | 4 |
| 7 | 778.3 | 389.7 | 761.3 | 760.3 | E | 375.2 | | 358.2 | 357.2 | 3 |
| 8 | 849.3 | 425.2 | 832.3 | 831.3 | A | 246.2 | | 229.1 | | 2 |
| 9 | 1,023.4 | 512.2 | 1,006.4 | 1,005.4 | R | 175.1 | | 158.1 | | 1 |

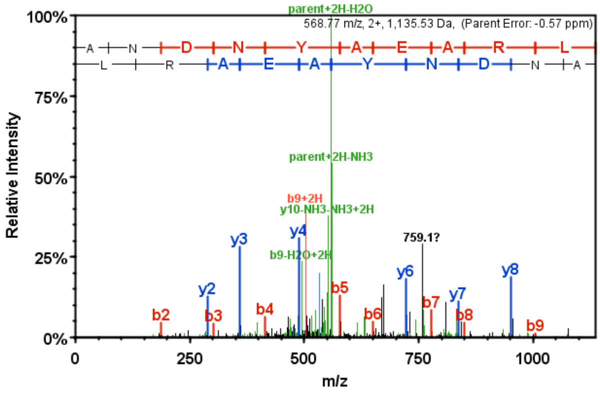


| B | B Ions | B+2H | B-NH3 | B-H2O | AA | Y Ions | Y+2H | Y-NH3 | Y-H2O | Y |
|---|---|---|---|---|---|---|---|---|---|---|
| 1 | 72.0 | | | | A | 1,136.5 | 568.8 | 1,119.5 | 1,118.5 | 10 |
| 2 | 186.1 | | 169.1 | | N | 1,065.5 | 533.3 | 1,048.5 | 1,047.5 | 9 |
| 3 | 301.1 | | 284.1 | 283.1 | D | 951.5 | 476.2 | 934.4 | 933.4 | 8 |
| 4 | 415.2 | | 398.1 | 397.1 | N | 836.4 | 418.7 | 819.4 | 818.4 | 7 |
| 5 | 578.2 | | 561.2 | 560.2 | Y | 722.4 | 361.7 | 705.4 | 704.4 | 6 |
| 6 | 649.3 | 325.1 | 632.2 | 631.2 | A | 559.3 | | 542.3 | 541.3 | 5 |
| 7 | 778.3 | 389.7 | 761.3 | 760.3 | E | 488.3 | | 471.3 | 470.3 | 4 |
| 8 | 849.3 | 425.2 | 832.3 | 831.3 | A | 359.2 | | 342.2 | | 3 |
| 9 | 1,005.4 | 503.2 | 988.4 | 987.4 | R | 288.2 | | 271.2 | | 2 |
| 10 | 1,136.5 | 568.8 | 1,119.5 | 1,118.5 | L | 132.1 | | | | 1 |

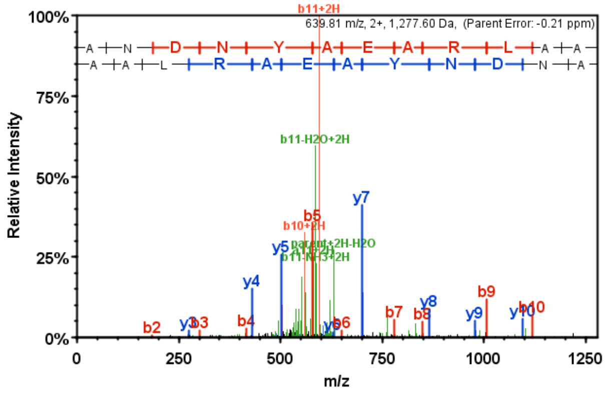


| B | B Ions | B+2H | B-NH3 | B-H2O | AA | Y Ions | Y+2H | Y-NH3 | Y-H2O | Y |
|---|---|---|---|---|---|---|---|---|---|---|
| 1 | 72.0 | | | | A | 1,278.6 | 639.8 | 1,261.6 | 1,260.6 | 12 |
| 2 | 186.1 | | 169.1 | | N | 1,207.6 | 604.3 | 1,190.5 | 1,189.6 | 11 |
| 3 | 301.1 | | 284.1 | 283.1 | D | 1,093.5 | 547.3 | 1,076.5 | 1,075.5 | 10 |
| 4 | 415.2 | | 398.1 | 397.1 | N | 978.5 | 489.8 | 961.5 | 960.5 | 9 |
| 5 | 578.2 | | 561.2 | 560.2 | Y | 864.5 | 432.7 | 847.4 | 846.4 | 8 |
| 6 | 649.3 | 325.1 | 632.2 | 631.2 | A | 701.4 | 351.2 | 684.4 | 683.4 | 7 |
| 7 | 778.3 | 389.7 | 761.3 | 760.3 | E | 630.4 | 315.7 | 613.3 | 612.3 | 6 |
| 8 | 849.3 | 425.2 | 832.3 | 831.3 | A | 501.3 | | 484.3 | 483.3 | 5 |
| 9 | 1,005.4 | 503.2 | 988.4 | 987.4 | R | 430.3 | | 413.3 | | 4 |
| 10 | 1,118.5 | 559.8 | 1,101.5 | 1,100.5 | L | 274.2 | | | | 3 |
| 11 | 1,189.6 | 595.3 | 1,172.5 | 1,171.5 | A | 161.1 | | | | 2 |
| 12 | 1,278.6 | 639.8 | 1,261.6 | 1,260.6 | A | 90.1 | | | | 1 |

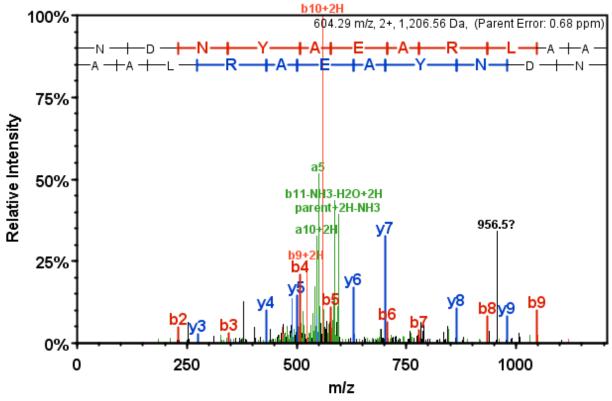


| B | B Ions | B+2H | B-NH3 | B-H2O | AA | Y Ions | Y+2H | Y-NH3 | Y-H2O | Y |
|---|---|---|---|---|---|---|---|---|---|---|
| 1 | 115.1 | | 98.0 | | N | 1,207.6 | 604.3 | 1,190.5 | 1,189.6 | 11 |
| 2 | 230.1 | | 213.1 | 212.1 | D | 1,093.5 | 547.3 | 1,076.5 | 1,075.5 | 10 |
| 3 | 344.1 | | 327.1 | 326.1 | N | 978.5 | 489.8 | 961.5 | 960.5 | 9 |
| 4 | 507.2 | | 490.2 | 489.2 | Y | 864.5 | 432.7 | 847.4 | 846.4 | 8 |
| 5 | 578.2 | | 561.2 | 560.2 | A | 701.4 | 351.2 | 684.4 | 683.4 | 7 |
| 6 | 707.3 | 354.1 | 690.2 | 689.3 | E | 630.4 | 315.7 | 613.3 | 612.3 | 6 |
| 7 | 778.3 | 389.7 | 761.3 | 760.3 | A | 501.3 | | 484.3 | 483.3 | 5 |
| 8 | 934.4 | 467.7 | 917.4 | 916.4 | R | 430.3 | | 413.3 | 412.3 | 4 |
| 9 | 1,047.5 | 524.2 | 1,030.5 | 1,029.5 | L | 274.2 | | | | 3 |
| 10 | 1,118.5 | 559.8 | 1,101.5 | 1,100.5 | A | 161.1 | | | | 2 |
| 11 | 1,207.6 | 604.3 | 1,190.5 | 1,189.6 | A | 90.1 | | | | 1 |

See next page

**Fig. S5B (continuation)**

**Spectra to Fig. 5C. SEP identifier in Table S4: CP004140.1:2290880-2290951:+; tmRNA-encoded proteolytic tag**

1 Peptide of CP004140.1:2290880-2290951:+ (Sequence matching one of the above 4 peptides),
3 PSMs with standard iPtgxDB in MM after SPE-enrichment and Lys-C digest

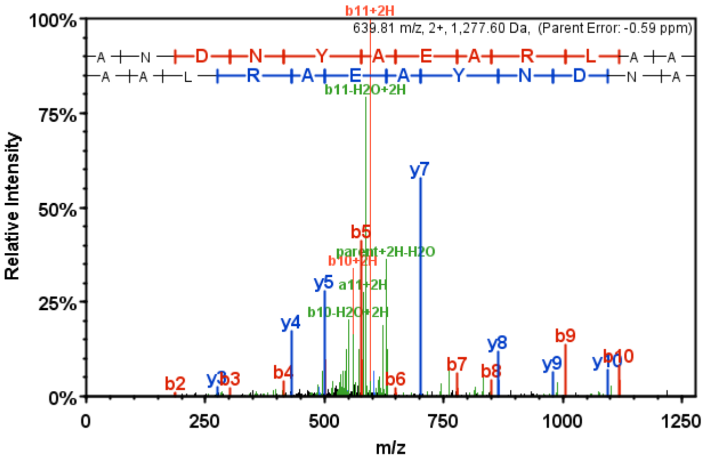


| B | B Ions | B+2H | B-NH3 | B-H2O | AA | Y Ions | Y+2H | Y-NH3 | Y-H2O | Y |
|---|---|---|---|---|---|---|---|---|---|---|
| 1 | 72.0 | | | | A | 1,278.6 | 639.8 | 1,261.6 | 1,260.6 | 12 |
| 2 | 186.1 | | 169.1 | | N | 1,207.6 | 604.3 | 1,190.5 | 1,189.6 | 11 |
| 3 | 301.1 | | 284.1 | 283.1 | D | 1,093.5 | 547.3 | 1,076.5 | 1,075.5 | 10 |
| 4 | 415.2 | | 398.1 | 397.1 | N | 978.5 | 489.8 | 961.5 | 960.5 | 9 |
| 5 | 578.2 | | 561.2 | 560.2 | Y | 864.5 | 432.7 | 847.4 | 846.4 | 8 |
| 6 | 649.3 | 325.1 | 632.2 | 631.2 | A | 701.4 | 351.2 | 684.4 | 683.4 | 7 |
| 7 | 778.3 | 389.7 | 761.3 | 760.3 | E | 630.4 | 315.7 | 613.3 | 612.3 | 6 |
| 8 | 849.3 | 425.2 | 832.3 | 831.3 | A | 501.3 | | 484.3 | 483.3 | 5 |
| 9 | 1,005.4 | 503.2 | 988.4 | 987.4 | R | 430.3 | | 413.3 | | 4 |
| 10 | 1,118.5 | 559.8 | 1,101.5 | 1,100.5 | L | 274.2 | | | | 3 |
| 11 | 1,189.6 | 595.3 | 1,172.5 | 1,171.5 | A | 161.1 | | | | 2 |
| 12 | 1,278.6 | 639.8 | 1,261.6 | 1,260.6 | A | 90.1 | | | | 1 |

**Fig. S5C**

**Spectra to Fig. 5C. SEP identifier in Table S4: CP004139.1:340914-341018:+; internal (nested) sORF**

3 PSM with standard iPtgxDB in MM after tryptic digest of non-enriched proteins

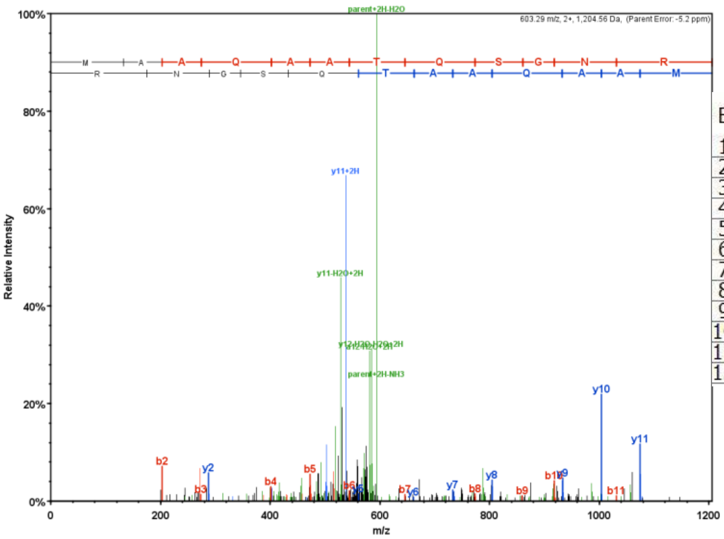


| B | B Ions | B+2H | B-NH3 | B-H2O | AA | Y Ions | Y+2H | Y-NH3 | Y-H2O | Y |
|---|---|---|---|---|---|---|---|---|---|---|
| 1 | 132.0 | | | | M | 1,205.6 | 603.3 | 1,188.5 | 1,187.6 | 12 |
| 2 | 203.1 | | | | A | 1,074.5 | 537.8 | 1,057.5 | 1,056.5 | 11 |
| 3 | 274.1 | | | | A | 1,003.5 | 502.2 | 986.5 | 985.5 | 10 |
| 4 | 402.2 | | 385.2 | | Q | 932.5 | 466.7 | 915.4 | 914.4 | 9 |
| 5 | 473.2 | | 456.2 | | A | 804.4 | 402.7 | 787.4 | 786.4 | 8 |
| 6 | 544.3 | 272.6 | 527.2 | | A | 733.4 | 367.2 | 716.3 | 715.3 | 7 |
| 7 | 645.3 | 323.2 | 628.3 | 627.3 | I | 662.3 | 331.7 | 645.3 | 644.3 | 6 |
| 8 | 773.4 | 387.2 | 756.3 | 755.4 | Q | 561.3 | | 544.2 | 543.3 | 5 |
| 9 | 860.4 | 430.7 | 843.4 | 842.4 | S | 433.2 | | 416.2 | 415.2 | 4 |
| 10 | 917.4 | 459.2 | 900.4 | 899.4 | G | 346.2 | | 329.2 | | 3 |
| 11 | 1,031.5 | 516.2 | 1,014.4 | 1,013.4 | N | 289.2 | | 272.1 | | 2 |
| 12 | 1,205.6 | 603.3 | 1,188.5 | 1,187.6 | R | 175.1 | | 158.1 | | 1 |

See next page

**Fig. S5D**

**Spectra of SEP7, identifier in Table S4: CP004140.1:2181026-2181205:+**

1 Peptide, 2 PSM with standard and custom iPtgxDB in MM after tryptic digest of non-enriched proteins

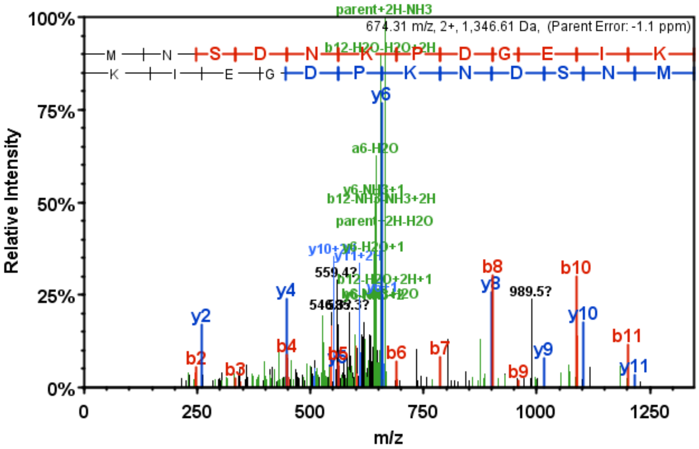


| B | B Ions | B+2H | B-NH3 | B-H2O | AA | Y Ions | Y+2H | Y-NH3 | Y-H2O | Y |
|---|---|---|---|---|---|---|---|---|---|---|
| 1 | 132.0 | | | | M | 1,347.6 | 674.3 | 1,330.6 | 1,329.6 | 12 |
| 2 | 246.1 | | 229.1 | | N | 1,216.6 | 608.8 | 1,199.6 | 1,198.6 | 11 |
| 3 | 333.1 | | 316.1 | 315.1 | S | 1,102.5 | 551.8 | 1,085.5 | 1,084.5 | 10 |
| 4 | 448.1 | | 431.1 | 430.1 | D | 1,015.5 | 508.3 | 998.5 | 997.5 | 9 |
| 5 | 562.2 | | 545.2 | 544.2 | N | 900.5 | 450.7 | 883.5 | 882.5 | 8 |
| 6 | 690.3 | 345.6 | 673.3 | 672.3 | K | 786.4 | 393.7 | 769.4 | 768.4 | 7 |
| 7 | 787.3 | 394.2 | 770.3 | 769.3 | P | 658.3 | 329.7 | 641.3 | 640.3 | 6 |
| 8 | 902.4 | 451.7 | 885.3 | 884.4 | D | 561.3 | | 544.3 | 543.3 | 5 |
| 9 | 959.4 | 480.2 | 942.4 | 941.4 | G | 446.3 | | 429.2 | 428.3 | 4 |
| 10 | 1,088.4 | 544.7 | 1,071.4 | 1,070.4 | E | 389.2 | | 372.2 | 371.2 | 3 |
| 11 | 1,201.5 | 601.3 | 1,184.5 | 1,183.5 | I | 260.2 | | 243.2 | | 2 |
| 12 | 1,347.6 | 674.3 | 1,330.6 | 1,329.6 | K | 147.1 | | 130.1 | | 1 |

**Fig. S5E**

**Spectrum of SEP1, identifier in Table S4: CP004140.1:1048709-1048780:+**

1 PSM with custom iPtgxDB in MM after SPE-enrichment, without proteolytic digest

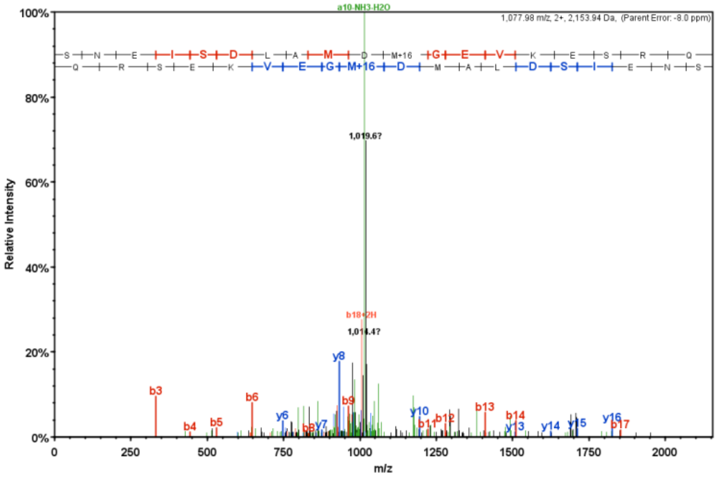


| B | B Ions | B+2H | B-NH3 | B-H2O | AA | Y Ions | Y+2H | Y-NH3 | Y-H2O | Y |
|---|---|---|---|---|---|---|---|---|---|---|
| 1 | 88.0 | | | 70.0 | S | 2,155.0 | 1,078.0 | 2,137.9 | 2,137.0 | 19 |
| 2 | 202.1 | | 185.1 | 184.1 | N | 2,067.9 | 1,034.5 | 2,050.9 | 2,049.9 | 18 |
| 3 | 331.1 | | 314.1 | 313.1 | E | 1,953.9 | 977.4 | 1,936.9 | 1,935.9 | 17 |
| 4 | 444.2 | | 427.2 | 426.2 | I | 1,824.8 | 912.9 | 1,807.8 | 1,806.8 | 16 |
| 5 | 531.2 | | 514.2 | 513.2 | S | 1,711.8 | 856.4 | 1,694.7 | 1,693.8 | 15 |
| 6 | 646.3 | 323.6 | 629.2 | 628.3 | D | 1,624.7 | 812.9 | 1,607.7 | 1,606.7 | 14 |
| 7 | 759.4 | 380.2 | 742.3 | 741.3 | L | 1,509.7 | 755.4 | 1,492.7 | 1,491.7 | 13 |
| 8 | 830.4 | 415.7 | 813.4 | 812.4 | A | 1,396.6 | 698.8 | 1,379.6 | 1,378.6 | 12 |
| 9 | 961.4 | 481.2 | 944.4 | 943.4 | M | 1,325.6 | 663.3 | 1,308.6 | 1,307.6 | 11 |
| 10 | 1,076.5 | 538.7 | 1,059.4 | 1,058.4 | D | 1,194.5 | 597.8 | 1,177.5 | 1,176.5 | 10 |
| 11 | 1,223.5 | 612.2 | 1,206.5 | 1,205.5 | M+16 | 1,079.5 | 540.3 | 1,062.5 | 1,061.5 | 9 |
| 12 | 1,280.5 | 640.8 | 1,263.5 | 1,262.5 | G | 932.5 | 466.7 | 915.5 | 914.5 | 8 |
| 13 | 1,409.6 | 705.3 | 1,392.5 | 1,391.5 | E | 875.5 | 438.2 | 858.4 | 857.4 | 7 |
| 14 | 1,508.6 | 754.8 | 1,491.6 | 1,490.6 | V | 746.4 | 373.7 | 729.4 | 728.4 | 6 |
| 15 | 1,636.7 | 818.9 | 1,619.7 | 1,618.7 | K | 647.3 | 324.2 | 630.3 | 629.3 | 5 |
| 16 | 1,765.8 | 883.4 | 1,748.7 | 1,747.8 | E | 519.3 | | 502.2 | 501.2 | 4 |
| 17 | 1,852.8 | 926.9 | 1,835.8 | 1,834.8 | S | 390.2 | | 373.2 | 372.2 | 3 |
| 18 | 2,008.9 | 1,005.0 | 1,991.9 | 1,990.9 | R | 303.2 | | 286.2 | | 2 |
| 19 | 2,155.0 | 1,078.0 | 2,137.9 | 2,137.0 | Q | 147.1 | | 130.0 | | 1 |

See next page

**Fig. S5F**
**Spectrum of SEP20, identifier in Table S4: CP004140.1:995681-995821:+**

1 PSM with custom iPtgxDB in MM after tryptic digest of non-enriched proteins

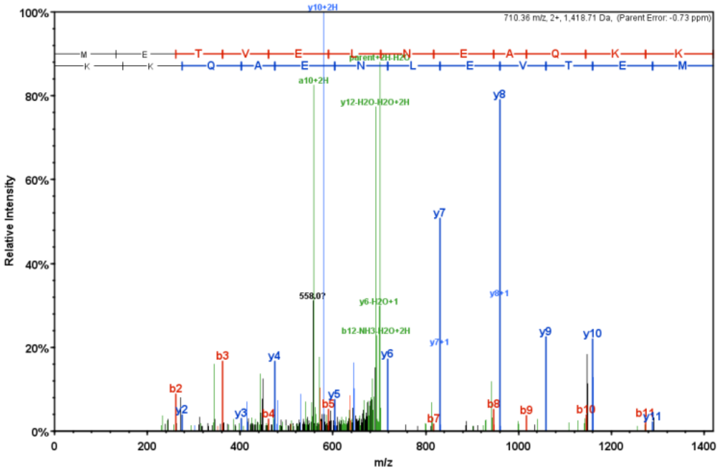


| B | B Ions | B+2H | B-NH3 | B-H2O | AA | Y Ions | Y+2H | Y-NH3 | Y-H2O | Y |
|---|---|---|---|---|---|---|---|---|---|---|
| 1 | 132.0 | | | | M | 1,419.7 | 710.4 | 1,402.7 | 1,401.7 | 12 |
| 2 | 261.1 | | | 243.1 | E | 1,288.7 | 644.8 | 1,271.6 | 1,270.7 | 11 |
| 3 | 362.1 | | | 344.1 | T | 1,159.6 | 580.3 | 1,142.6 | 1,141.6 | 10 |
| 4 | 461.2 | | | 443.2 | V | 1,058.6 | 529.8 | 1,041.6 | 1,040.6 | 9 |
| 5 | 590.2 | | | 572.2 | E | 959.5 | 480.3 | 942.5 | 941.5 | 8 |
| 6 | 703.3 | 352.2 | | 685.3 | L | 830.5 | 415.7 | 813.4 | 812.5 | 7 |
| 7 | 817.4 | 409.2 | 800.3 | 799.4 | N | 717.4 | 359.2 | 700.4 | 699.4 | 6 |
| 8 | 946.4 | 473.7 | 929.4 | 928.4 | E | 603.3 | 302.2 | 586.3 | 585.3 | 5 |
| 9 | 1,017.5 | 509.2 | 1,000.4 | 999.4 | A | 474.3 | 237.7 | 457.3 | | 4 |
| 10 | 1,145.5 | 573.3 | 1,128.5 | 1,127.5 | Q | 403.3 | 202.1 | 386.2 | | 3 |
| 11 | 1,273.6 | 637.3 | 1,256.6 | 1,255.6 | K | 275.2 | 138.1 | 258.2 | | 2 |
| 12 | 1,419.7 | 710.4 | 1,402.7 | 1,401.7 | K | 147.1 | | 130.1 | | 1 |

**Fig. S5G**
**Spectrum of a highly conserved SEP (64 aa), identifier in Table S4: CP004139.1:748169-748363:-**

3 PSM with standard iPtgxDB in TY after SPE-enrichment, without proteolytic digest

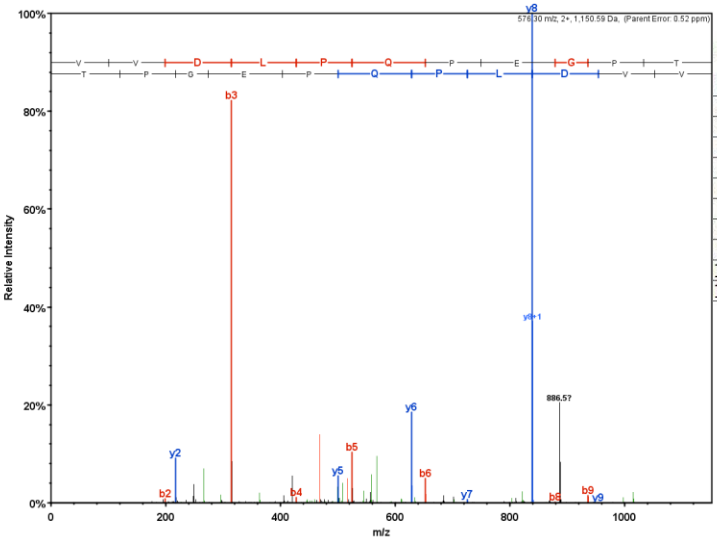


| B | B Ions | B+2H | B-NH3 | B-H2O | AA | Y Ions | Y+2H | Y-NH3 | Y-H2O | Y |
|---|---|---|---|---|---|---|---|---|---|---|
| 1 | 100.1 | | | | V | 1,151.6 | | 1,134.6 | 1,133.6 | 11 |
| 2 | 199.1 | | | | V | 1,052.5 | | 1,035.5 | 1,034.5 | 10 |
| 3 | 314.2 | | | 296.2 | D | 953.5 | | 936.4 | 935.4 | 9 |
| 4 | 427.3 | | | 409.2 | L | 838.4 | | 821.4 | 820.4 | 8 |
| 5 | 524.3 | | | 506.3 | P | 725.3 | | 708.3 | 707.3 | 7 |
| 6 | 652.4 | 326.7 | 635.3 | 634.4 | Q | 628.3 | | 611.3 | 610.3 | 6 |
| 7 | 749.4 | 375.2 | 732.4 | 731.4 | P | 500.2 | | | 482.2 | 5 |
| 8 | 878.5 | 439.7 | 861.4 | 860.5 | E | 403.2 | | | 385.2 | 4 |
| 9 | 935.5 | 468.2 | 918.5 | 917.5 | G | 274.1 | | | 256.1 | 3 |
| 10 | 1,032.5 | 516.8 | 1,015.5 | 1,014.5 | P | 217.1 | | | 199.1 | 2 |
| 11 | 1,151.6 | 576.3 | 1,134.6 | 1,133.6 | I | 120.1 | | | 102.1 | 1 |

**Figure S5. Mass spectrometry of selected novel small open reading frame-encoded proteins (SEPs).** Here, we show some of the spectra that allowed us to identify novel SEPs. If more than one peptide spectrum match (PSM) was detected for a given peptide ion, a representative spectrum was selected. MS2 spectra with assigned fragment ion m/z (left) and fragmentation tables (right) were obtained with Scaffold V4.8.7 using the search output files (*.sf3), which were deposited at the ProteomeXchange Consortium with the dataset identifier PXD034931. Colored m/z were assigned in the identifying MS2 spectra.

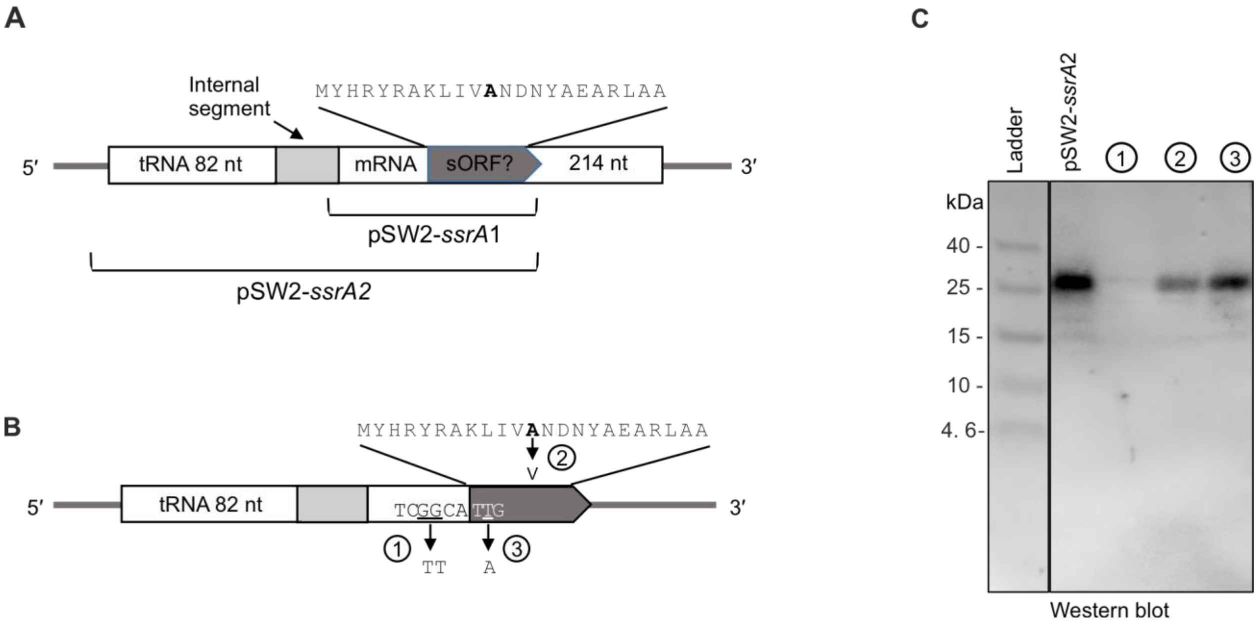


**Figure S6. Analysis of a putative small open reading frame (sORF) in tmRNA of *Sinorhizobium meliloti* 2011. (A)** Schematic view of the chromosomal *ssrA* locus corresponding to tmRNA, which is discontinuous in Alphaproteobacteria due to post-transcriptional removal of the indicated internal segment (Keiler et al. 2000; Ulvé et al. 2007). The tRNA and mRNA parts of the tmRNA and their lengths are indicated. In the mRNA part, a putative sORF corresponding to a 23-amino acid (aa) SEP was predicted (the potential SEP sequence is shown). The 3′ part of the sORF corresponds to the proteolytic tag-encoding sequence (the alanine encoded by the resume codon is shown in bold). The indicated parts of the *ssrA* gene were cloned in pSW2, and the *S. meliloti* 2011 strains containing the corresponding plasmids were used for Western blot analysis. While no specific bands were detected in lysates of the pSW2-ssrA1-containing plasmid, which lacks the tRNA part of tmRNA (data not shown), a strong signal was obtained with the pSW2-ssrA2 plasmid, which contains both the tRNA and mRNA parts (see panel C). **(B)** Schematic representation of the *ssrA* part cloned in pSW2-ssrA2. The used mutations are indicated. 1: Conserved GG nucleotides upstream of the putative start codon TTG were mutated to TT. 2: The resume codon was changed to encode valine instead of alanine. 3: The putative start codon TTG was mutated to the stop codon TAG. **(C)** Western blot analysis with antibodies directed against the FLAG part of the sequential peptide affinity (SPA) tag, which was fused in frame to the proteolytic tag. Crude lysates (corresponding to 20 OD) of *S. meliloti* 2011 strains containing the indicated plasmids (see panels A and B) were analyzed. The detected bands above 25 kDA have identical lengths. Expression of the corresponding SPA-tagged peptide was abolished by the indicated GG/TT mutation, which disrupts conserved base pairing in the tmRNA (Keiler et al. 2000), whereas a weak signal was still detected when the resume codon was mutated. Destroying the putative 23 aa sORF (TTG/TAG mutation in variant 3) did not abolish the expression of the tagged peptide. These results, combined with no detection of a peptide using pSW2-ssrA1, suggest that the SPA-tagged peptide detected in panel C corresponds to the 12 aa proteolytic tag and not to the putative 23 aa SEP. Furthermore, the data support the important role of the analyzed GG nucleotides for the tmRNA function.

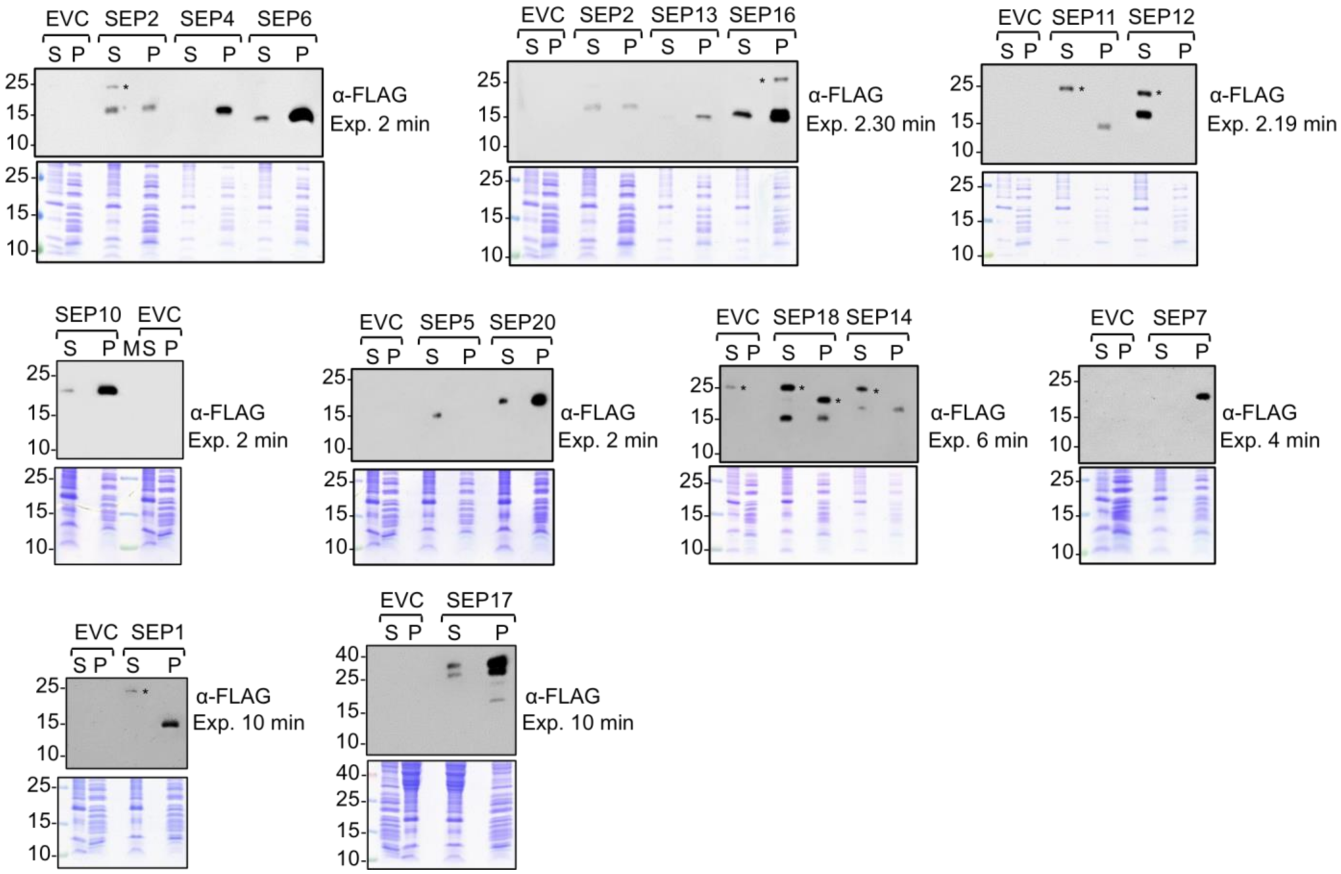


**Figure S7. Analysis of the S100 and P100 fractions of *Sinorhizobium meliloti* 2011 strains producing the indicated small open reading frame-encoded proteins (SEPs) from pSW2-SEP plasmids.** The empty vector control (EVC) strain was used as the negative control. Identical volumes of the S100 and P100 fractions were loaded. Top panels: Western blot analysis with monoclonal anti-FLAG antibodies. Bottom panels: Coomassie-stained gels, in which the used protein fractions are shown. Migration of marker proteins (in kDa) is shown on the left side, and exposition times are indicated. * Unspecific signal.
